# Rewiring of chromatin regulation underlies the evolution of brown algal multicellularity

**DOI:** 10.1101/2025.09.16.676480

**Authors:** Jeromine Vigneau, Jaruwatana Sodai Lotharukpong, Pengfei Liu, Remy Luthringer, Bérangère Lombard, Damarys Loew, Fabian B. Haas, Michael Borg, Susana M Coelho

## Abstract

Chromatin structure plays a central role in regulating transcription, genome stability, and epigenetic inheritance in eukaryotes. Much of our understanding of chromatin architecture and histone post-translational modifications (hPTMs) comes from a narrow set of animal and plant models, but emerging data from non-model lineages are challenging canonical views of how chromatin functions across the tree of life. Brown algae are complex multicellular eukaryotes that provide a unique perspective on chromatin evolution given their independent origin of complex multicellularity. Here, we compile the chromatin toolkit of brown algae and show that canonical silencing systems involving DNA cytosine methylation and PRC2-mediated H3K27 methylation were lost early in their evolution. By generating hPTM profiles from diverse brown algal clades, we resolve the nature and regulatory roles of chromatin states in this lineage and show how H3K79 methylation emerged and diversified as a repressive system. We further uncover sex-specific reconfigurations in species with varying degrees of sexual dimorphism and reconstruct the ancestral regulatory landscape that likely preceded the emergence of brown algae. Together, our findings illuminate the dynamic evolution of chromatin regulation in a distinct multicellular lineage and challenge assumptions about the universality of chromatin-based mechanisms across eukaryotes.

## Introduction

In eukaryotes, chromatin regulates access to the genome by packaging DNA into an organised structure that modulates transcription and other DNA-based processes. This organization is defined by chromatin states, which consist of specific combinations of DNA and/or histone post-translational modifications (hPTMs), DNA-binding proteins, and 3D structural features^1–3^. Chromatin states are often dynamic and responsive to developmental and environmental cues, and is some cases can be stably maintained though epigenetic inheritance^4^. In addition to gene regulation, chromatin plays a critical role in safeguarding genome integrity by repressing transposable elements and other invasive mobile elements^5^.

Most of our understanding of epigenetic and chromatin-based regulation has largely been informed by the discovery and analysis of hPTMs in a limited range of plant and animal models^6^. While many hPTMs are evolutionarily conserved across eukaryotic lineages and trace back to the last eukaryotic common ancestor (LECA), recent work on non-model lineages is revealing a surprising diversity of chromatin systems and challenges classical views about the conserved function of hPTMs^7^. For example, the brown alga *Ectocarpus* lacks canonical hPTMs such as H3K9 and H3K27 methylation, while H3K79 methylation appears to play a repressive role rather than associating with active transcription as in yeast and metazoans, highlighting a striking divergence in chromatin regulation^8–11^. The brown algae represent a distinct branch of complex multicellular eukaryotes, separated from animals and plants, that have evolved within the last 450 million years and display a broad diversity in genome size, morphology, and sexual systems^12–14^. With the recent development of genomic tools for this clade, including over 65 genome assemblies of which several are at a chromosome scale, as well as genetic tools, brown algae have emerged as attractive model organisms for comparative studies. Notably, their genomes have remained largely syntenic^15^, which facilitates cross-species comparative genomic studies.

Here, we investigated the diversity of chromatin landscapes across major brown algal lineages that encompasses the broad morphological complexity and variation in sexual systems within the group, along with an outgroup species for comparison. We combined genome-wide chromatin modification maps with gene expression data and various genomic features to functionally interpret chromatin states across brown algae. Our analyses reveal several key insights into the evolution of chromatin landscapes in this group. First, the emergence of the brown algal lineage was accompanied by the loss of both PRC2-mediated repression and DNA methylation, representing a major shift in epigenetic regulation during early brown algal evolution. Second, activation-associated hPTMs are highly conserved across species, suggesting functional constraint. In contrast, the repressive role of H3K79 methylation does not appear to be ancestral but likely evolved prior to the divergence of the Ectocarpales and Laminariales. Third, sex-specific chromatin state differences between male and female gametophytes seems independent from the degree of sexual dimorphism observed at the organismal level, suggesting that morphological sex differentiation may be driven by chromatin reconfiguration at a limited number of high-order effector loci. Finally, by examining chromatin and DNA methylation in an outgroup species, we trace the transition from an ancestral epigenetic landscape to the distinct architecture observed in brown algae.

## Results

### Evolution of chromatin and epigenetic-related genes

The recent availability of several high-quality brown algal genomes allowed us to investigate the conservation and evolution of chromatin-associated proteins across the clade^12,15^. Using BLAST searches and orthology-based approaches, we screened for homologs of known chromatin-related proteins and revealed a conserved and distinct repertoire across brown algae (**Fig. 1A**). Consistent with previous reports, we observed a complete absence of MET1 orthologs and other DNA methyltransferases across the clade. MET1 orthologs were only identified in the closest non-Phaeophycean relative *Schizocladia ischiensis*, suggesting that the lineage-specific loss of DNA methyltransferases occurred early in brown algal evolution. Similarly, we found that EED and SUZ12 subunits specific to the Polycomb Repressive Complex 2 (PRC2) are absent from all surveyed brown algal species, while the modular MSI1 subunit common to other chromatin complexes was present. EED and SUZ12 orthologs are also absent in *S. ischiensis* as well as other closely-related Ochrophytina species, suggesting that the loss of PRC2 may have preceded the emergence of the Phaeophyceae. The loss of PRC2 was also reflected in the absence of PRC1 homologs, which forms a distinct repressive complex in animals and plants^16^. SET domain proteins are abundant across the clade, which included homologs of Trithorax-related histone methyltransferases and two brown algal-specific families of SET domain proteins. Interestingly, orthologs of DOT1, which is responsible for histone H3 lysine 79 (H3K79) methylation in yeast and animals^17^, are greatly expanded among brown algae and have diversified into at least five distinct families (**Fig. S1**), highlighting a Phaeophyceae-specific adaptation in chromatin regulation. These observations show that the emergence of the brown algal lineage is marked by the concurrent loss of epigenetic control via PRC2 and DNA methylation and an expansion of DOT1 histone methyltransferases, suggesting a major shift in gene regulatory strategies in this clade.

**Figure 1.**
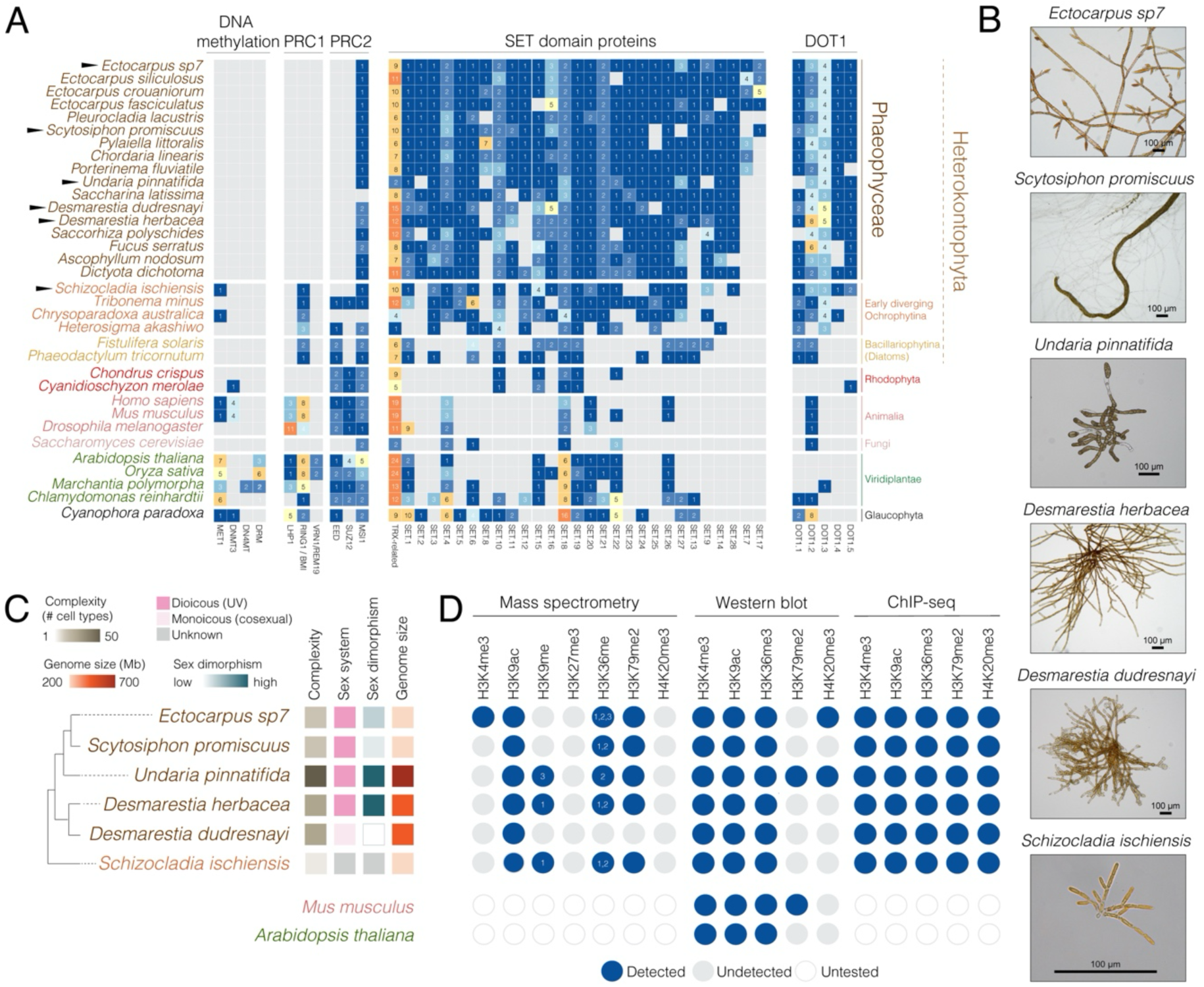
Evolution of chromatin-related and epigenetic pathways in brown algae. (**A**) Census of chromatin proteins across the brown algae, their closest relatives, and other major eukaryotic lineages. The species focused on in this study are indicated by a black triangle. (**B**) Representative images of the species used in this study. (**C**) Schematic diagram summarising the phylogenetic relationship of the six species studied alongside biological features that distinguish them, namely their overall morphological complexity, sexual system, sexual dimorphism and relative genome size. (**D**) Summary of the major hPTMs detected in the panel of six species using mass spectrometry, western blot and/or ChIP-seq. *Ectocarpus* mass spectrometry data was produced in a previous study^8^.

To investigate the diversity of chromatin landscapes in brown algae, we selected a set of representative species spanning the phylogenetic breadth of the group, chosen to reflect variation in morphological complexity, sexual systems, and reproductive strategies (**Fig. 1B-C**). To enable comparisons between male and female gametophytes within the same genetic background, we used sibling samples where possible, thereby minimizing genetic variability unrelated to sex (**Table S1**). Our sampling included *Desmarestia dudresnayi* that recently transitioned to co-sexuality (monoicy), alongside its closest relative *Desmarestia herbacea,* a dioicous species with separate male and female individuals. We also included *Undaria pinnatifida*, a kelp species characterized by a highly complex morphology, high sexual dimorphism, and a large genome with expanded UV sex chromosomes^15^. This contrasts with *Ectocarpus* sp.7 and *Scytosciphon promiscuus*, which possess smaller sex-linked regions on their UV sex chromosomes and have low-to-medium sexual dimorphism, respectively^15,18,19^ . Finally, we included the filamentous chromista *S. ischiensis* as a representative outgroup species from the closest diverging lineage outside of the brown algae^20^.

We first performed mass spectrometry of histone preparations to detect hPTMs across the five representative species, as done previously in *Ectocarpus*^9^ (**Fig. 1D; Table S2**). Overall, hPTMs were detected with minimal species-specific differences. Histones that we identified close to known H2A.Z variants (H2A.Z-like) carry a heavily acetylated tails are present in *Ectocarpus*, *U. pinnatifida*, *D. herbacea* and *S. ischiensis* but not retrieved in *S. promiscuus* and *D. dudresnayi*. H3K9me1 was detected in *D. herbacea* and *S. ischiensis*, and H3K9me3 in *U. pinnatifida*, but neither mark was observed in the other species (**Fig. 1D; Table S2**). Importantly, methylated forms of histone H3K27 were absent from all five brown algal species analysed, consistent with the lack of PRC2 subunits encoded in their genomes (**Fig. 1A, D; Table S2**). Taken together, this data indicates that the lack of canonical epigenetic regulation via DNA and H3K27 methylation is a general feature of brown algae.

In earlier studies focusing on chromatin landscapes in the filamentous brown alga *Ectocarpus*, we identified H3K4me3, H3K9ac and H3K36me3 as hPTMs of active chromatin, whereas H3K79me2 and H4K20me3 were more strongly linked to transcriptional repression ^8,9^. Aside for H3K4me3 and H4K20me3, which were technically challenging to isolate in our mass spectrometry runs, we were able to detect most of the other hPTMs in each species (**Table S2**). We further validated the presence of H3K4me3, H3K9ac and H3K36me3 in each species using immunoblotting although H3K79me2 was only detectable in *U. pinnatifida* (**Fig. 1D; Fig. S2** for full length uncropped blots). We speculate that H3K79me2 is likely present at low levels and is challenging to detect with immunoblotting, which was further supported by similarly low signals in mouse histone extracts (**Fig. S2**). Building on their conservation across the clade and prior characterisation in *Ectocarpus*, we focused on this set of five hPTMs to investigate the evolutionary dynamics of chromatin landscapes in brown algae.

### The chromatin landscape of diverse brown macroalgae

Closely related male and female gametophyte (haploid) lines were used to generate sex-specific ChIP-seq profiles for the five histone PTMs in species with separate sexes (*Ectocarpus*, *S. promiscuus*, *U. pinnatifida*, *D. herbacea*) (**Table S1**; see Materials and Methods). Monoicous *D. dudrenayi* gametophytes and vegetative tissue from *S. ischiensis* were similarly profiled (**Fig. 2A**, **Fig. S3A**). For each species, we profiled at least 400 clonal individuals per male, female, monoicous or vegetative replicate, then merged ChIP-seq replicates after confirming high reproducibility (**Fig. S4; Table S4**).

**Figure 2.**
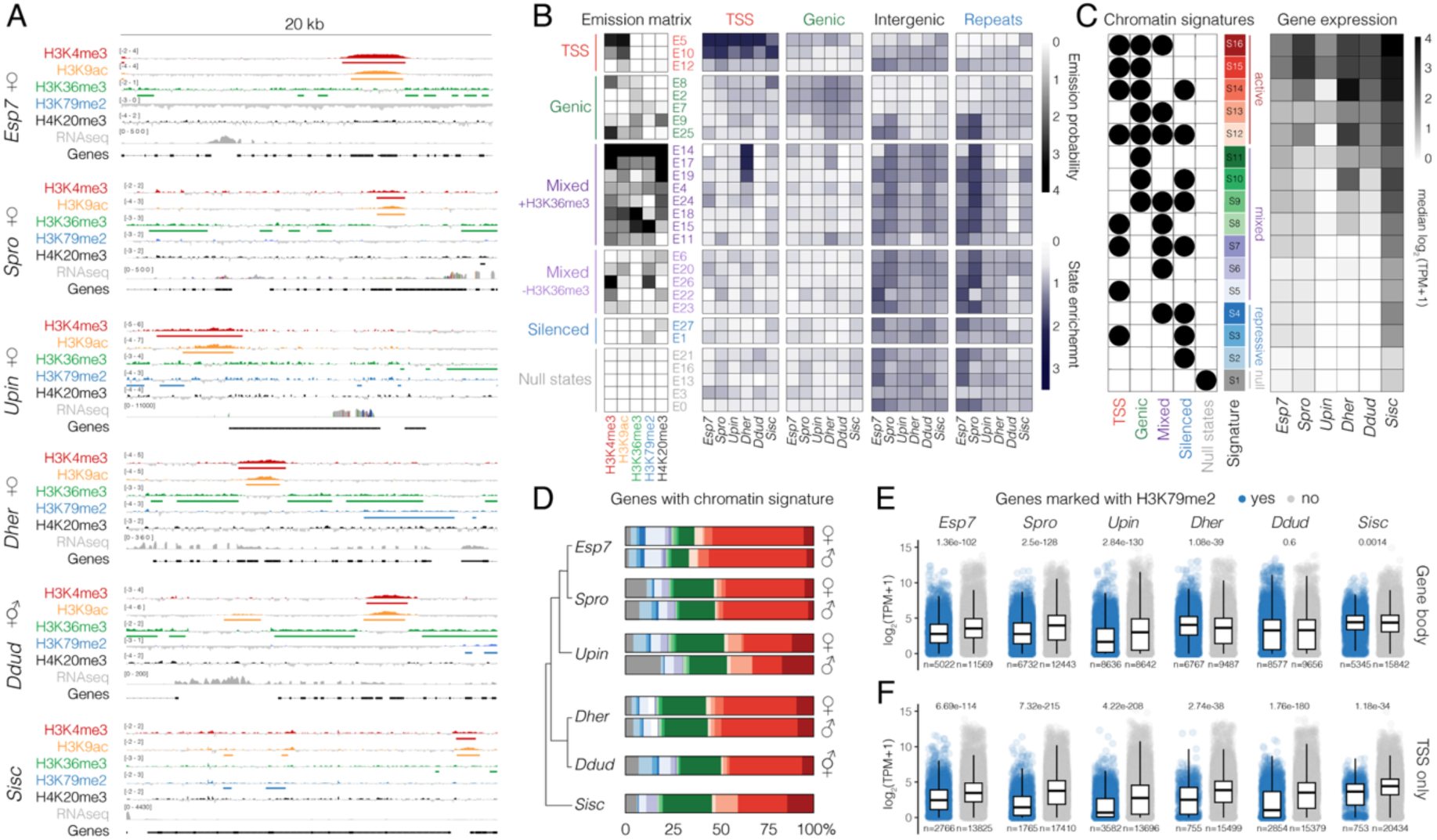
The chromatin landscape across multiple brown algae. (**A**) Representative genome browser tracks of ChIP-seq and RNA-seq datasets in each species. Data for female gametophyte is shown for dioicous species. ChIP-seq coverage is represented as the log_2_ ratio of IP DNA relative to histone H3, with the range indicated on each track. ChIP-seq peaks for each hPTM are indicated under their respective track (see **Fig. S4A** for male data). (B) A model of chromatin emission states inferred by hiHMM across all ChIP-seq datasets (left) alongside the enrichment of each chromatin state in genomic features of each species (right). Female samples are shown for dioicous species (see **Fig. S4B** for male data). (**C**) Matrix of the chromatin signatures assigned to genes based on the hiHMM emission state model (see methods) summarising the proportion of repressive, mixed, and active chromatin states within each signature. The right panel displays the median RNA-seq expression level (log_2_ TPM+1) of genes associated with each signature. Female samples are shown for dioicous specie s (see **Fig. S4C** for male data). (**D**) Proportion of chromatin signatures assigned to genes in each sample. (**E-F**) Gene expression with (blue) or without (grey) an H3K79me2 peak overlapping their gene body (**E**) or TSS (**F**) in each species. Female samples are shown for dioicous species (see **Fig. S4D-E** for male data). *P*-values were computed via the Kruskal–Wallis test in (**E**-**F**).

Metaplot analyses revealed that H3K4me3 and H3K9ac were highly enriched at transcription start sites (TSSs) and positively correlated with gene expression across all brown algal species (**Fig. 2A; Fig. S5**). H3K36me3 was consistently enriched over the body of expressed genes and showed a positive correlation with transcript levels, although this varied and was most pronounced in the two Ectocarpales species. In contrast, H3K79me2 was broadly deposited across gene bodies but showed a mark depletion at the most highly expressed genes. In *Ectocarpus*, *S. promiscuus*, *U. pinnatifida*, elevated H3K79me2 levels were strongly associated with reduced gene expression, whereas this correlation was restricted to the TSS in the two *Desmarestia* species. While the outgroup species *S. ischiensis* showed similar hPTM deposition patterns, enrichment signals and overall correlations were weaker, with the strongest association observed for H3K9ac at the TSS and the weakest for H3K79me2. These findings indicate that hPTMs associated with active transcription function similarly as in other eukaryotic lineages, while the deposition pattern of H3K79me2 and its association with transcriptionally repressed genes differs across brown algae.

Next, we employed the Bayesian non-parametric framework hiHMM to jointly infer chromatin state maps across the six different species^21^ (**Suppl. Dataset 1**). This analysis produced a probabilistic model composed of 27 emission states (**Fig. 2B, Fig. S6**), which we grouped into five broad categories based on their enrichment in specific genomic features: three TSS-associated states, five genic states, 14 mixed states distinguished by the presence or absence of H3K36me3, two silenced states, and five ‘null’ states lacking any of the assayed hPTMs (**Fig. 2B**, **Fig. S4B**). Closer inspection revealed both conserved and species-specific patterns of chromatin state occurrence (**Fig. S7**). For example, TSSs in all species were defined by a highly conserved and limited combination of states enriched for H3K4me3 and H3K9ac (E5, E10, E12). Gene bodies were predominantly characterised by three genic states common to all six species (E8, E2, E7). In contrast, intergenic regions and repeats exhibited more variable patterns that included a combination of mixed and silenced states. Notably, two genic states (E9, E25) were exclusively enriched in intergenic regions and repeats in *Ectocarpus* and *S. promiscuus*, suggesting a more derived chromatin landscape in the Ectocarpales lineage.

Multiple emission states can occur along the length of a single gene in various combinations, making it challenging to analyse chromatin state dynamics at the gene level. To address this, we applied our previously established approach to determine the presence of the five broad chromatin state categories at each gene across all species, resulting in 16 distinct combinations of chromatin signatures (see Materials and Methods section) (**Fig. 2C, Fig. S3C, Table S5-10**). Based on the predominant hPTMs associated with each chromatin state, we further classified the chromatin signatures into four main groups: active (S12-S16), mixed (S5-S11), repressive (S2-S4) and null (S1) (**Fig. 2C, Fig. S4C**). The relative distribution of chromatin signatures assigned to genes was broadly conserved among species, with active signatures S15 and S16 representing the most common categories (**Fig. 2D**). We verified the relationship between the 16 chromatin signatures and gene expression in each species using paired RNA-seq data generated from the same biological material used for ChIP-seq profiling (**Fig. 2C**, **Fig. S4C**, **Fig. S8, Fig. S9**, **Table S4**). Across all brown algal species, genes assigned to active signatures consistently had higher transcript levels than those with mixed signatures, while genes associated with repressive or null signatures showed the lowest levels overall (**Fig. 2C; Fig. S9**). In contrast, the outgroup *S. ischiensis* exhibited a much weaker correlation between chromatin signatures and gene expression, particularly for repressive signatures, suggesting a distinct regulatory system.

To better understand the relationship between H3K79me2 and transcriptional repression, we compared expression of genes marked with and without H3K79me2 in each species. In *Ectocarpus*, *S. promiscuus* and *U. pinnatifida*, H3K79me2-marked genes had significantly lower expression than unmarked genes (**Fig. 2E, Fig. S4D**). This repression was most pronounced in *U. pinnatifida*, where genes assigned chromatin signatures S12 and S14 containing H3K79me2 were largely silenced relative to their equivalent signatures S15 and S16 without H3K79me2 (**Fig. 2C**, **Fig. S4C**). In contrast, H3K79me2-marked genes in the two *Desmarestia* species did not show reduced expression compared to unmarked genes (**Fig. 2E**, **Fig. S4D**). However, metaplot analyses suggested that the association of H3K79me2 with repression may be spatially restricted to the TSS in this lineage (**Fig. S4**). Indeed, parsing genes by the presence or absence of H3K79me2 at the TSS recapitulated the repression patterns observed in the other brown algae (**Fig. 2F**, **Fig. S4E**). TSS-restricted silencing by H3K79me2 was also evident in the *U. pinnatifida* male, which also happened to have a higher proportion of null signature S1 compared to the female and other species (**Fig. 2D**). This was likely due to a high abundance of sperm cells in the *U. pinnatifida* male tissue we processed, leading us to focus the remainder of our cross-species comparisons using female samples. In the outgroup *S. ischiensis*, TSS-associated H3K79me2 also correlated with reduced gene expression, although this was limited to a relatively small subset of genes (**Fig. 2F**). These results suggest that the repressive role of H3K79me2 arose early in the brown algal lineage that has since undergone further functional diversification over the course of their evolution.

### Conserved function of chromatin signatures among brown algae

The brown algae are thought to have emerged around 450 million years ago, making them one of the youngest complex multicellular lineages, a fact that is reflected by the strong synteny observed across their genomes^12,22^. This prompted us to examine the evolutionary dynamics of chromatin signatures across the six species. For this, we first identified 3143 conserved single copy orthologs and assessed whether they retained similar signatures, using female data for species with separate sexes (**Table S11**)^8^. Our analysis revealed a strong overall conservation of chromatin signatures, which significantly exceeded expectation based of permutation tests assuming no conservation (**Fig. 3A**). Pairwise comparisons with model alga *Ectocarpus* further showed that the conservation of chromatin signatures correlated with phylogenetic distance (**Fig. 3B**). In *Ectocarpus*, these single copy orthologs were strongly enriched for the transcriptionally active signature S15 and largely corresponded to deeply conserved genes with essential cellular functions and broad expression patterns, suggesting that they primarily represent housekeeping genes (**Fig. S10**). These results highlight how orthologous genes have maintained similar active chromatin organisation across brown algal evolution, which is likely driven by their constitutive expression and core housekeeping function.

**Figure 3.**
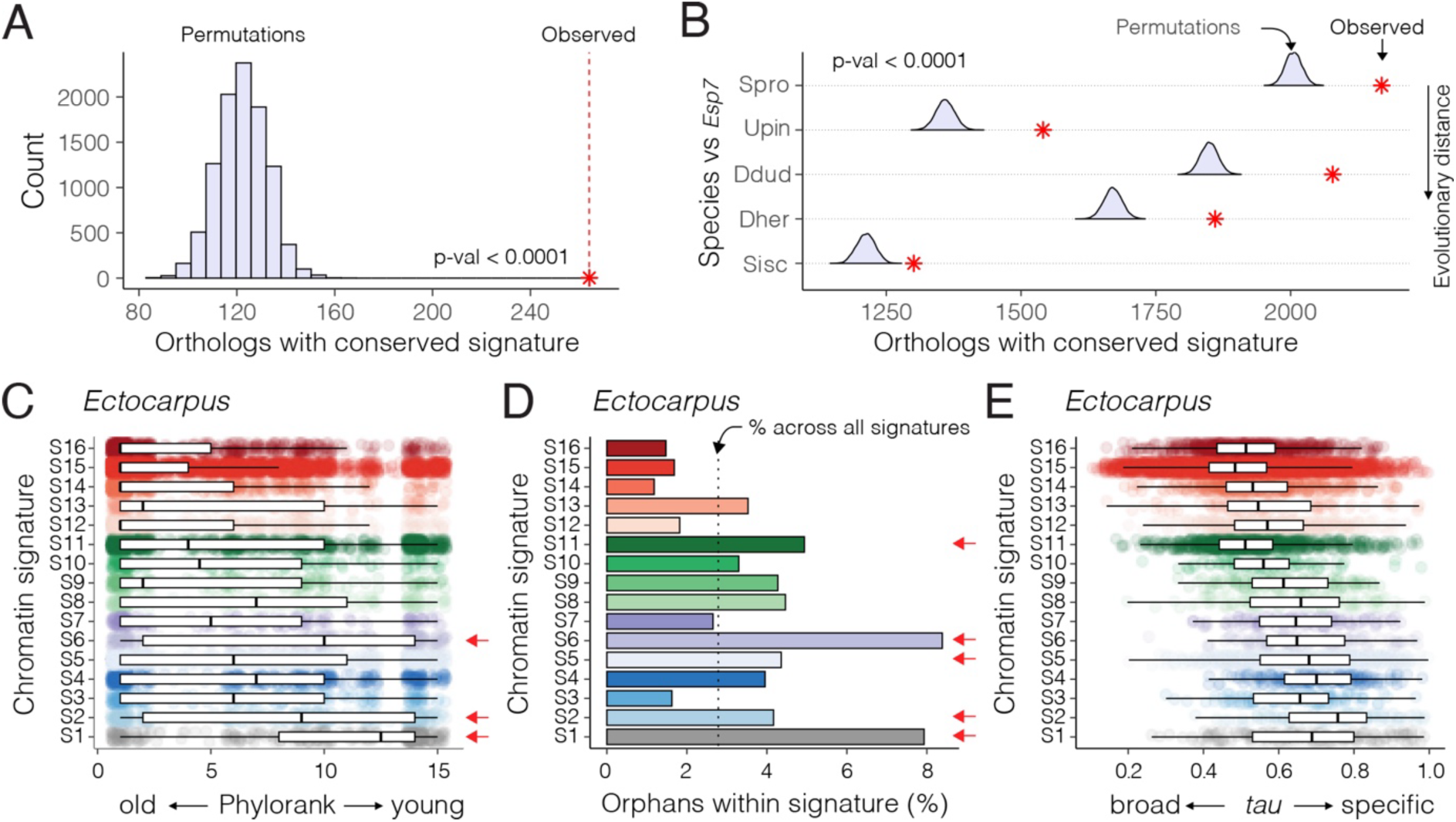
Conservation and phylostratigraphy of chromatin signatures across brown algae. (**A**) Conservation of chromatin signatures at single copy orthologs across all six species tested alongside permutations assuming no conservation. (**B**) Conservation of chromatin signatures in pairwise comparisons with the model alga *Ectocarpus* alongside permutations assuming no conservation. *P*-values were computed from the 10,000 permutation results. (**C**) Distribution of gene age across each chromatin signature in *Ectocarpus*. (**D**) Percentage of orphan genes assigned to each chromatin signature in *Ectocarpus*. (**E**) Distribution of expression specificity scores (*tau*) for genes assigned to each chromatin signature in *Ectocarpus*.

We next asked whether chromatin signatures were associated with genes of similar evolutionary age and expression profile ^23^. We used genomic phylostratigraphy to infer the evolutionary age of genes then assessed the distribution of their assigned signature, revealing statistically significant differences (**Fig. 3C**; **Fig. S11**; Kruskal–Wallis test; p < 2.2e-16 for all species). Notably, null and repressive chromatin signatures were more strongly associated with evolutionarily younger genes in *Ectocarpus* and across the other species (**Fig. 3C**; **Fig S11**). Moreover, when limiting our analysis to species-specific orphan genes, we found that these were strongly enriched in chromatin signatures with reduced expression (S1, S2, S5, S6, S11) both in *Ectocarpus* and across the clade (**Fig. 2C**; **Fig. 3D; Fig. S12**). This suggests that younger genes typically reside in heterochromatic regions when compared to older and more conserved genes. Differences in chromatin signatures also manifested in similar expression dynamics. Genes with null and repressive chromatin signatures exhibited more restricted expression profiles than those with active signatures, a pattern that was consistent across all species, indicating that these signatures may represent facultative heterochromatin potentially involved in developmental gene regulation (**Fig. 3E; Fig. S13**). Taken together, our findings demonstrate that chromatin signatures function similarly across 450 million years of brown algal evolution^13^, revealing conserved chromatin features underlying gene regulation in this lineage.

### The chromatin landscape involved in sex determination and differentiation

UV sex chromosomes have distinctive genomic and evolutionary features due to their mode of inheritance and their characteristic sex-determining regions (SDRs) that do not recombine^22,24,25^. In *Ectocarpus*, these features are reflected by a unique chromatin landscape on the UV chromosomes, where a much higher proportion of genes are marked by repressive chromatin signatures than on autosomes^8,10^. Strikingly, this pattern was highly conserved across all five brown algal species we examined, with sex chromosome genes consistently showing reduced proportions of active signatures and increased proportions of null and repressive signatures than autosomal genes (**Fig. 4A**). The UV sex chromosomes also consistently displayed a markedly lower proportion of conserved chromatin signatures than their autosomal counterparts (**Fig. 4B**). These findings show that the UV sex chromosomes have a distinct chromatin organization across brown algae, which reflects the rapid gene turnover that is characteristic for these fast-evolving chromosomes.

**Figure 4.**
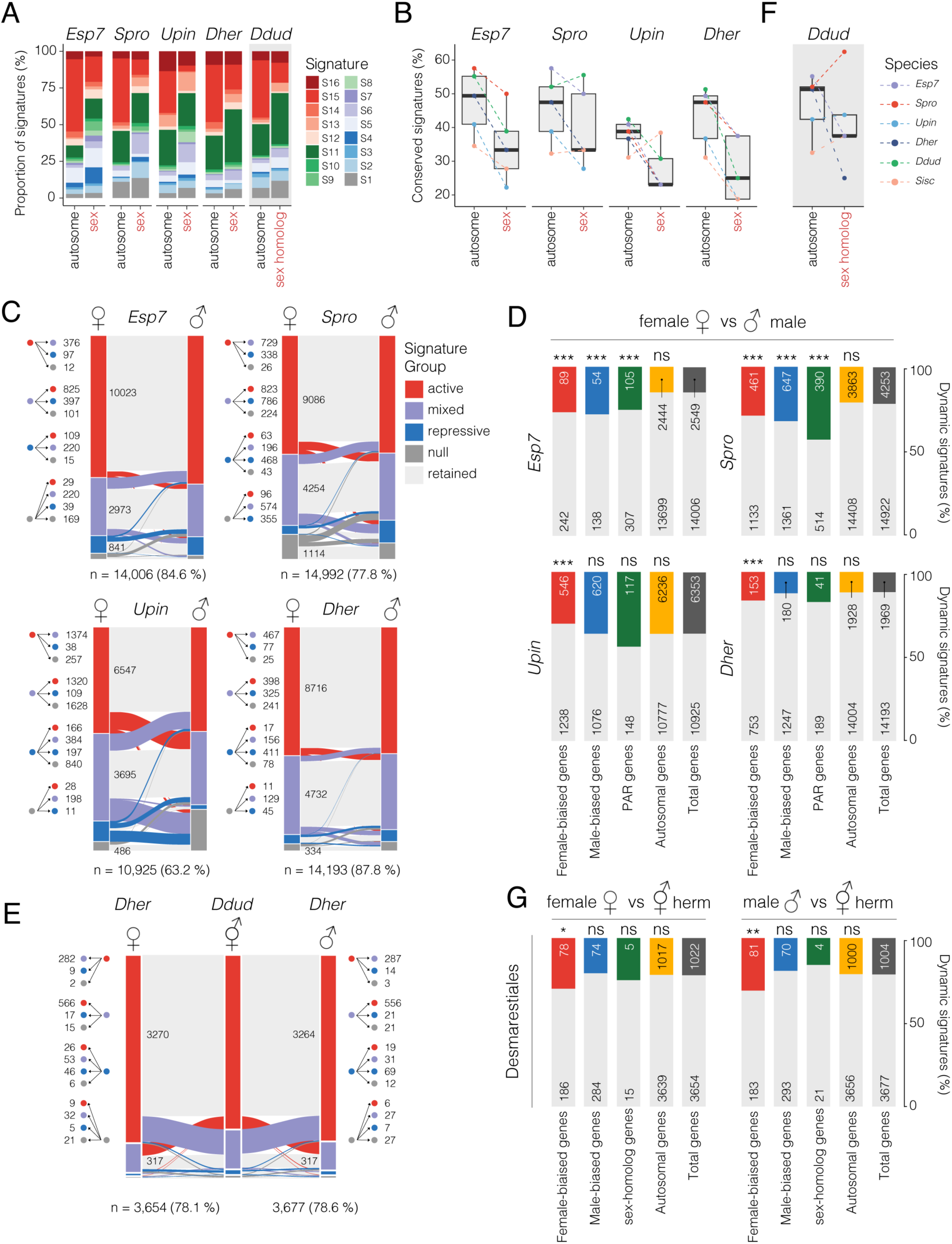
Chromatin reconfiguration during sexual differentiation in brown algae with contrasting sexual systems. (**A**) Proportion of chromatin signatures on autosomes versus sex chromosomes across dioicous brown algal species. (**B**) Chromatin signature conservation between autosomes and sex chromosomes in each species. (C) Major reconfiguration in chromatin signatures between females and males. The direction of chromatin signature changes and the number of genes involved are shown on the left , with the total number of genes shown below. (**D**) Proportion of sex-biased, PAR and autosomal genes that dynamically switch chromatin signature between sexes. *P*-values of a χ^2^ analysis on top of each bar indicate whether the proportions observed differ significantly from the genome average. (**E**) Major reconfiguration in chromatin signatures between females and males versus co-sexual *Desmarestia* species. The direction of chromatin signature changes and the number of genes involved are shown on the left, with the total number of genes shown below. (**F**) Chromatin signature conservation across all autosomes versus the sex homolog (ancestral sex chromosome) in the co-sexual species *Desmarestia dudresnayi*. (**G**) Proportion of sex-biased, PAR and autosomal genes that dynamically switch chromatin signatures in *D. herbacea* (dioicous) versus *D. dudresnayi* (monoicous).

We next assessed chromatin dynamics involved in sexual differentiation by comparing chromatin signatures between males and females. To capture these dynamics, we focused on major chromatin reconfigurations by comparing shifts between the four major chromatin signature groups (active, mixed, repressive and null). Across the four dioicous species, the vast majority of genes (63.2 - 87.8 %) retained the same chromatin signature group between sexes (**Fig. 4C**). Of those that were dynamic between sexes, sex-biased genes (i.e., genes that were differentially expressed between sexes) were significantly more likely to undergo major reconfiguration in *Ectocarpus* and *S. promiscuus* compared to the genomic background (**Fig. 4D**). In *D. herbacea and U. pinnatifida*, only female-biased genes were enriched for such changes (**Fig. 4D**). Genes located in the pseudoautosomal region (PAR) of the UV chromosomes were significantly enriched for dynamic chromatin reconfigurations in *Ectocarpus* and *S. promiscuus* (**Fig. 4D**). Taken together, these results suggest that major chromatin reconfigurations underlying sex-biased gene regulation are subtle and variable across brown algae and are largely restricted to specific loci, particularly female-biased genes.

Next, we focused on major chromatin reconfigurations associated with co-sexuality by comparing male and female gametophytes of the dioicous species *D. herbacea* with monoicous gametophytes of *D. dudresnayi*. Similar to the dioicous species, the vast majority of genes retained the same chromatin signature group between the co-sexual and either sex of *D. herbacea* (**Fig. 4E**). As observed during sexual differentiation in the dioicous species, major chromatin reconfigurations were preferentially associated with orthologous female-biased genes, whereas orthologous male-biased genes showed no dynamic differences in the co-sexuals compared to females, and were even less dynamic when compared to males (**Fig. 4E**). These results suggest that major chromatin reconfigurations underlying co-sexuality largely occur at female-biased genes, paralleling the situation observed in dioicy. Finally, we asked whether the transition to co-sexuality influences the conservation of chromatin signatures on the UV sex chromosomes by testing whether the reduced conservation underlying dioicy is retained when a former sex chromosome becomes an autosome (hereafter termed the ‘sex-homolog’^22^). Intriguingly, the *D. dudresnayi* sex-homolog also showed reduced conservation of chromatin signatures, indicating that this property of sex chromosomes persists after the transition to monoicy (**Fig. 4F**). Thus, the former *D. dudresnayi* sex chromosome retains molecular footprints of its past life as a sex chromosome and can leave lasting evolutionary imprints at the chromatin level.

### Epigenetic control in an ancestor of the brown algae

The outgroup *S. ischiensis* stood out among the six species we analysed due to its weaker correlation between chromatin signatures and gene expression, and its relatively low number of H3K79me2-marked genes. This prompted us to explore other potential regulatory mechanisms that may operate in *S. ischiensis*. Our phylogenetic analysis revealed that, unlike brown algae, *S. ischiensis* encodes a MET1 ortholog (**Fig. 1A**), leading us to speculate that DNA methylation may regulate gene expression in this species. To explore this, we called different forms of DNA methylation from Oxford Nanopore long-read sequences used to assemble the *S. ischiensis* genome. The level of 4-methylcytosine and 6-methyladenine calls are unlikely to be genuine in *S. ischiensis* since their genome-wide levels were very low (< 5%) and comparable to those detected on the plastid genome (**Fig. S14**). In contrast, 5-methylcytosine in a CG context (5mCG) was highly abundant, reaching almost 80% genome-wide and far exceeding background levels on the plastid genome (**Fig. 5A-B**). By comparison, 5mC in CHG and CHH contexts was much lower and showed no preference for TEs as they do in plants, suggesting that these methylation patterns are unlikely to have a specific function in *S. ischiensis* (**Fig. S14**).

**Figure 5.**
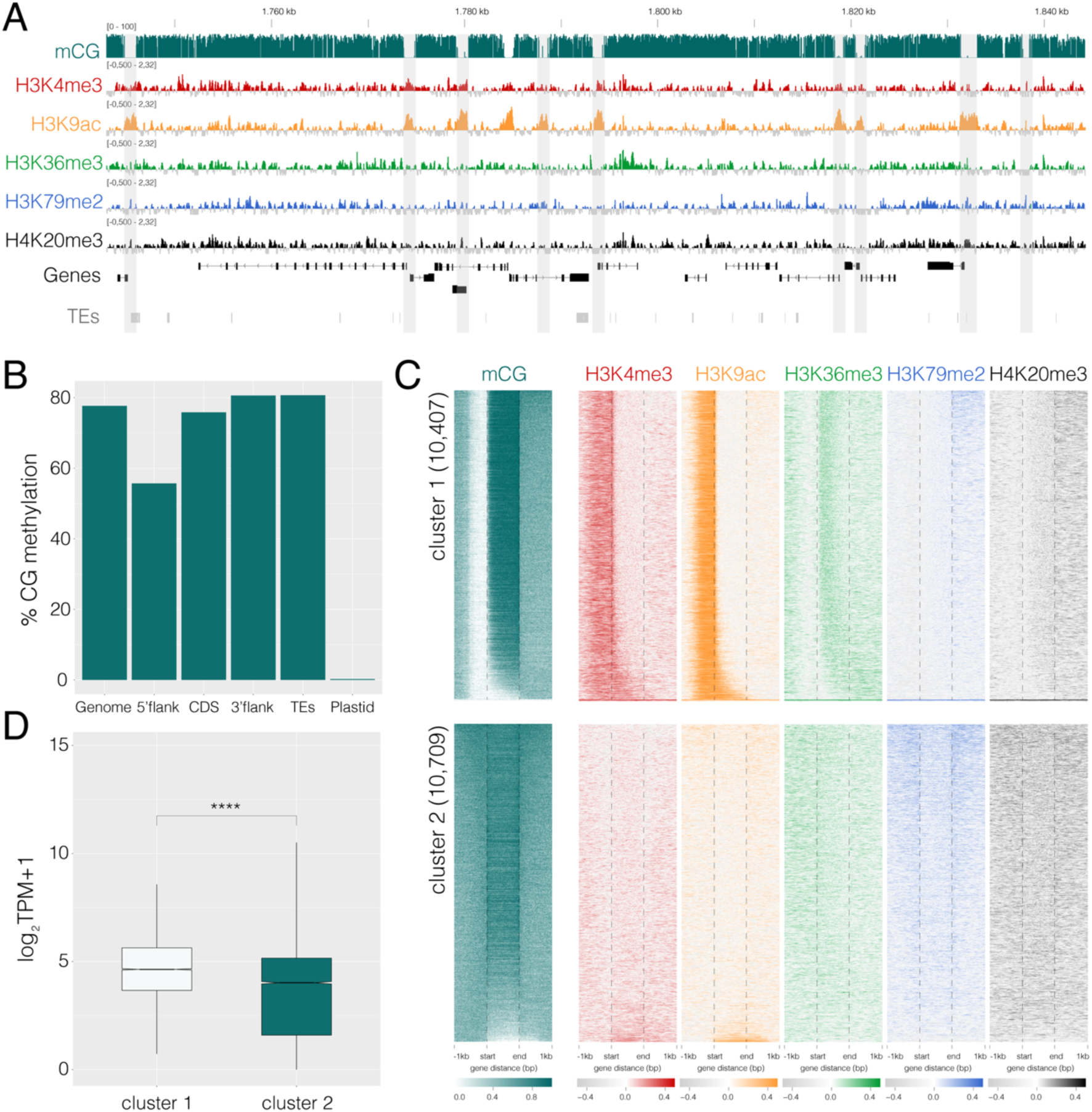
Epigenetic landscape in the closest outgroup of the brown algae. (**A**) Genome browser view of ChIP-seq and CG methylation tracks in *Schizocladia ischiensis*. Demethylated CpG-like islands are shown with grey shading. (**B**) Average genome-wide CG methylation levels at different genomic features of the nuclear genome and plastid genome. (**C**) Chromatin and CG methylation landscape over *S. ischiensis* genes clusters based on the presence (cluster 1) and absence (cluster 2) of CG methylation at promoter regions. (**D**) Gene expression level of genes forming part of cluster 1 and cluster 2 shown in panel C.

Closer inspection of 5mCG in *S. ischiensis* revealed discrete regions with a complete loss of methylation around the promoter of protein-coding genes (**Fig. 5A**), which is reminiscent of unmethylated CpG “islands” found in mammalian genomes^26,27^. These demethylated islands coincided with the enrichment of TSS-associated H3K4me3 and H3K9ac, with the upstream region of genes consistently having reduced 5mCG levels relative to other genomic regions (**Fig. 5A-B**). Hierarchical clustering of 5mCG levels alongside the six profiled hPTMs revealed two main clusters distinguished by demethylated promoter regions and the concurrent enrichment of H3K4me3 and H3K9ac (**Fig. 5C**). Genes in this “active” cluster had higher transcript levels compared to genes without promoter demethylation, consistent with the deposition of active hPTMs (**Fig. 5D**). These results explain the weaker correlation of chromatin signatures in *S. ischiensis* (see **Fig. 2C**) and suggest that active DNA demethylation at promoters may have played a key regulatory role prior to the loss of DNA methylation during early brown algal evolution.

## Discussion

Brown algae are the most recent lineage to have independently evolved complex multicellularity, making them an important case study to understand how regulatory mechanisms evolve during the emergence of organismal complexity. Our comparative analysis across the Phaeophyceae and their closest relatives reveals that this transition was accompanied by a fundamental shift in chromatin regulation, where canonical repressive pathways involving PRC2, H3K9 and DNA cytosine methylation are largely dispensable across the clade. Although the presence of DNA methylation is highly variable across eukaryotes^28^, H3K27 methylation is largely ubiquitous and essential in animals, fungi, land plants, and even closely related Stramenopiles^29,30^. In brown algae, the concurrent loss of DNA, H3K9 and H3K27 methylation is unprecedented and underscores the unique evolutionary trajectory of this lineage.

Our analysis of the outgroup *S. ischiensis* provides an important perspective on the ancestral regulatory landscape of the brown algae. We show that the presence of DNA methyltransferases is accompanied by high genome-wide levels of 5mCG methylation. Interestingly, we reveal a strong link between promoter demethylation and the accumulation of active histone marks at expressed genes, suggesting a regulatory model in which DNA methylation plays a central role in gene repression. The localised demethylation of promoter regions and the coordinated enrichment of active histone marks point to a regulatory system that is strikingly similar to vertebrates rather than to modern-day brown algae^26,27^. The absence of PRC2 orthologs in *S. ischiensis* suggests that H3K27me3-mediated regulation is likely to have been lost early in brown algal evolution. The eventual loss of both DNA methylation and PRC2 in the common ancestor of brown algae, which is exceptionally rare among multicellular eukaryotes, would have necessitated the emergence of alternative repressive systems, providing fertile ground for the adaptation of DOT1-mediated H3K79 methylation into a key repressive pathway.

The expansion and diversification of DOT1-like enzymes into lineage-specific families is a distinguishing feature of Phaeophyceae evolution. In yeast and animals, DOT1-dependent H3K79 methylation regulates diverse processes including gene expression, replication initiation, DNA damage response, microtubule reorganization and chromosome segregation^17^. DOT1 enzymes also promote heterochromatin formation by regulating pericentromeric transcription of satellite repeats, where bursts of transcription are required to establish and maintain long-term silencing^31,32^. We speculate that this role in heterochromatin formation could have been co-opted during brown algal evolution to give rise to its repressive role in the modern-day. H3K79 methylation has similarly been implicated in gene repression in other eukaryotic lineages^7^, underscoring a capacity for the DOT1 pathway to be independently recruited to regulate silencing during eukaryotic evolution.

By investigating chromatin landscapes across diverse brown algal species, we established a defined set of combinatorial chromatin states (signatures) that are highly predictive of gene expression across the clade. Active transcription was consistently associated with H3K4me3, H3K9ac, and H3K36me3, whereas H3K79me2 and H4K20me3 were associated with reduced transcription. The localisation of H3K79me2 and its perceived repressive role varied among the brown algal clades, further highlighting the dynamic nature of this pathway. In addition to transcriptional states, chromatin signatures were also strongly correlated with the evolutionary age of genes and the breadth of their expression. Notably, repressive chromatin signatures were highly enriched among young species-specific genes with restricted gene expression patterns, a pattern mirrored across a wide range of eukaryotes^33–35^. This supports the idea that heterochromatic regions may serve as a cradle for the emergence of novel genes^22^, with silencing followed by eventual reprogramming and expression in reproductive cell types providing a route for the evolution and selection of new gene functions^35–37^.

Our findings have also shed light on the regulation of sex determination and sex chromosome evolution in brown algae. Brown algal UV sex chromosomes consistently show enrichment of repressive chromatin and reduced conservation of chromatin signatures compared to autosomes. These features are consistent with suppressed recombination and gene turnover in the sex-determining regions^15,38–40^ which appear to leave a lasting epigenomic footprint. Interestingly, even after the transition from dioicy to co-sexuality in the Desmarestiales, the sex-homolog retains a distinct chromatin profile, suggesting that the epigenomic legacy of sex chromosome identity persists long after recombination resumes. Chromatin modifications thus not only reflect current transcriptional states but may also shape the long-term evolutionary trajectory of the sex-homolog during transitions towards co-sexuality by constraining or biasing subsequent regulatory evolution. This echoes findings from plants and animals where epigenetic silencing marks, dosage-compensation mechanisms and heterochromatin expansion continues to influence genome function long after sex chromosome turnovers or fusions^41–44^. Such epigenomic legacies may therefore represent a general principle in the evolution of sex determination systems. We also uncovered major chromatin reconfigurations associated with sexual differentiation across brown algae. *Ectocarpus* and *S. promiscuus*, both species with limited sexual dimorphism, showed the most pronounced localised chromatin reconfigurations at PAR genes and sex-biased genes. In other species, such reconfigurations consistently occurred at female-biased genes, highlighting augmentation of the chromatin reconfiguration on the female program as the principal driver of sexual differentiation in brown algae with stronger sexual dimorphism, including those with co-sexual systems.

In summary, our findings highlight evolutionary innovations in the chromatin toolkit that accompanied the emergence of complex multicellularity in brown algae, where the loss of canonical repression pathways and the rise of DOT1/H3K79 methylation established a novel regulatory system that now underpin development and reproduction in this vital and unique eukaryotic lineage.

## Methods

### Genome screening and orthology inference

We initially performed genome screening to identify the components of epigenetic regulation in brown algae using blastp^45^ (default parameters) by leveraging recently published genomic data of species in the brown algal lineage (Phaeophyceae) and early-diverging Ochrophytina^12,15^ along with representatives from across major eukaryotic groups. We selected five species covering the brown algal phylogenetic diversity and one outgroup species. Orthology inference was performed on these six species using OrthoFinder v2.5.5^46^.

### Biological material

Gametophytes of the five brown algal species and the outgroup *S. ischiensis* were cultivated in 90cm Petri dishes (Corning® Gosselin™ BH90B-102) containing at least 10 individuals, with Provasoli enriched seawater as described in ^19,47^. Fertile individuals were harvested with a 70µm strainer, then rinsed with seawater and dry with a paper towel for further processing. Light and temperature conditions were optimised for fertility, as described in **Table S1**.

### hPTM profiling

#### Histones extraction

Histones were extracted from 0.5g of frozen algae, pulverized in liquid nitrogen. The powder was then homogenised in 40mL of M1 buffer (10mM Na Phosphate Buffer pH7, 100mM NaCl, 1000mM Hexylene Glycol, 10mM b-mercaptoethanol, 1X cOmplete Proteinase inhibitor Cocktail (Roche). After filtering through 2 layers of Miracloth (Milipore #475855), each sample was centrifugated at 2000 × g for 10 minutes at 4 °C. The pellet was carefully resuspended in 80mL M2 buffer (10mM Na Phosphate Buffer pH7, 100mM NaCl, 10mM MgCl2, 1000mM Hexylene Glycol, 0.1% Triton X-100, 10mM b-mercaptoethanol, 1X cOmplete™, EDTA-free Protease Inhibitor Cocktail (Roche, #CO-RO) twice. This pellet was then incubated with Extraction buffer (1000mM CaCl2, 20mM Tris-HCl pH7.25, 1x cOmplete cOmplete™, EDTA-free Protease Inhibitor Cocktail) for 10 minutes on ice. 0.3N of 37% HCl was added thereafter followed by centrifugation at 10 000 × g for 5 minutes at 4 °C. The resulting supernatant was collected in a fresh Protein LoBind® Tubes Eppendorf tubes. Histones were precipitated with 20% tricholoacetic acid (TCA) and incubated for 10 minutes before 13 000 × g centrifugation at 4 °C for 30 minutes, followed by successive washes with 20% ice-cold TCA, ice-cold acetone supplemented with 0.2% HCl and ice-cold acetone. The pellet was dried at room temperature and resuspended in miliQ water overnight at 4 °C.

#### Western blot

Histone samples were supplemented with Laemmli 2X and 100mM of DTT and NaOC until blue coloration was observed and incubated at 95 °C for 5 minutes. Histone PTMs were detected on a 15% handcast SDS-PAGE gel, using the same antibodies listed belowy as in the ChIP experiment. For H3, 3–15 µg of tissue-equivalent sample was loaded onto the gel. Histone samples, corresponding to 3–15 µg of tissue equivalent for H3, were loaded. For H3K9me3, around 10 ug of tissue-equivalent sample twice that amount was used, and for other histone marks, approximately 40 ug of tissue-equivalent sample four times or more. Proteins were transferred onto a 0.45µm nitrocellulose membrane (0.45µm, BioRad, #1620113) on a Trans-Blot® Turbo™ Transfer System (BioRad, #1704150). Membranes were blocked in 5% milk in 1× PBS-T for 30 minutes. Primary antibodies were diluted in 5% milk in 1× PBS-T and incubated for 1 hour at room temperature. These included rabbit anti-H3 (Histone H3 (D2B12) XP Rabbit mAb (ChIP Formulated), Cell Signaling Technology #4620S), anti-H3K4me3 (Tri-Methyl-Histone H3 (Lys4) (C42D8) Rabbit mAb, CST #9751S) , anti-H3K9ac (Acetyl-Histone H3 (Lys9) (C5B11) Rabbit mAb, CST, #9649S), anti-H3K79me2 (Di-Methyl-Histone H3 (Lys79) (D15E8) XP Rabbit mAb, CST, #5427S), and anti-H4K20me3 (Tri-Methyl-Histone H4 (Lys20) (D84D2) Rabbit mAb, CST, #5737S) at 1:1000 dilution, and anti-H3K36me3 (AbcamTri-Methyl-Histone H3 (Lys36) (D5A7) XP Rabbit mAb, Abcam, #4909S) at 1 µg/mL. Membranes were incubated with HRP-conjugated anti-rabbit secondary antibody (1:2000, CST #7074S). After further washes, membranes were developed using a 1:1 mix of Trans-Blot® Turbo™ Transfer System (BioRad, #1704150). Images were captured with ChemiDoc Imaging System from BioRad.

#### Mass spectrometry

Samples for liquid chromatography-tandem mass spectrometry (LC-MS/MS) were prepared by migrating the extracted histone on a 14% SDS-polyacryamide gel at 100V for 10 minutes. The gel was dyed with LabSafe GEL Blue™ (G-BIOSCIENCES, #786-35) following the manufacturer instruction. This was followed by LC-MS/MS, performed by coupling a Vanquish Neo LC system (Thermo Scientific) to an Orbitrap Astral mass spectrometer, interfaced by a Nanospray Flex ion source (Thermo Scientific). In a subsequent round of analyses, a RSLCnano system (Ultimate 3000, Thermo Scientific) to an Orbitrap Exploris 480 mass spectrometer (Thermo Scientific) was additionnally employed.

On the Vanquish Neo LC system, peptides were injected onto a C18 column (75 µm inner diameter x 50 cm double nanoViper PepMap Neo, 2μm, 100Å, Thermo Scientific) regulated also at 50 °C, and separated with a linear gradient from 100% buffer A’ to 28% buffer B at a flow rate of 300 nL/min over 104 minutes. The Orbitrap Astral mass spectrometer was run in Data Dependent Acquisition (DDA) mode and MS full scans were performed in the ultrahigh-field Orbitrap mass analyzer in ranges m/z 380–1200 (resolution of 240 000 at m/z 200; maximum injection time 100 ms; AGC 300%). For the Astral MS/MS spectra, the top N most intense ions were isolated and subjected to further fragmentation via high energy collision dissociation (HCD) activation with the auto gain control (AGC) target set to 100%. We selected ions with charge state from 2+ to 6+ for screening. Normalised collision energy (NCE) was set at 30 and the dynamic exclusion at 20s.

On the RSLCnano system, peptides were trapped on a C18 column (75 μm inner diameter × 2 cm; nanoViper Acclaim PepMapTM 100, Thermo Scientific) with buffer A (2/98 CH3CN/H2O in 0.1% formic acid) at a flow rate of 2.5 µL/min over 4 minutes. Separation was performed on a 50 cm × 75 μm C18 column (nanoViper Acclaim PepMapTM RSLC, 2 μm, 100Å, Thermo Scientific) regulated to a temperature of 50 °C with a linear gradient of 2% to 30% buffer B (100% CH3CN in 0.1% formic acid) at a flow rate of 300 nL/min over 91 minutes. On the Orbitrap Exploris 480 mass spectrometer, full scans were performed in ranges m/z 375–1500 (resolution of 120 000 at m/z 200; maximum injection time 25 ms; AGC 300%) and the top 20 most intense ions were isolated and subjected to further fragmentation via HCD activation at resolution of 15 000 with the AGC target set also to 100%. We also selected ions with charge state from 2+ to 6+. NCE was set at 30 and with a dynamic exclusion of 10s.

The resulting LC-MS/MS data was searched against the species-specific histone sequences using Mascot^48^. Enzyme specificity was set to trypsin and a maximum of five-missed cleavage sites were allowed. Oxidized methionine, carbamidomethylated cysteine, N-terminal acetylation, acetylation, methylation (mono, di and tri) of lysine, methylation (mono and di) of arginine, methylation of glutamic acid and aspartic acid were set as variable modifications and with a maximum of nine modifications for all Mascot searches. Maximum allowed mass deviation was set to 10 ppm for monoisotopic precursor ions and 0.02 Da for MS/MS peaks. The resulting Mascot files were further processed using myProMS (v.3.10; https://github.com/bioinfo-pf-curie/myproms)^49^.

### ChIP-seq

To map hPTMs to the genome, we performed ChIP-seq to detect the enrichment of H3K4me3, H3K9ac, H3K36me3, H3K79me2, and H4K20me3. Each sample was prepared from approximately 0.6 g of semi-dry algal tissue (∼600 individuals), which was then fixed in seawater containing 1% freshly prepared formaldehyde for 10 minutes. The fixed sample was quenched with 400 mM glycine in 1× PBS, followed by rinsing with fresh seawater to remove residual formaldehyde. Nuclei were isolated by grinding the cross-linked tissue in liquid nitrogen and resuspending the powder in a nuclear isolation buffer containing 0.1% Triton X-100, 125 mM sorbitol, 20 mM potassium citrate, 30 mM MgCl₂, 5 mM EDTA, 5 mM β-mercaptoethanol, 55 mM HEPES (pH 7.5), and 1× EDTA-free protease inhibitor cocktail (Roche #CO-RO). The suspension was homogenized using a Tenbroeck Potter, filtered through Miracloth (Millipore #475855), and centrifuged at 3000 × g for 10 minutes at 4 °C. The nuclear pellet was washed twice with the same buffer and once more with buffer lacking Triton X-100, conserving the centrifuge parameters. Nuclear pellets were then lysed in 1ml of nuclear lysis buffer total (1% SDS, 10 mM EDTA, 50 mM Tris-HCl pH 8, and protease inhibitors). Chromatin was fragmented via sonication using a Covaris E220 Evolution sonicator (settings: 25% duty cycle, 75 peak power, 200 cycles/burst, 900 s duration at 4 °C) in 8 microTUBE AFA Fiber Snap-Cap tubes. Cellular debris was cleared by centrifugation at 14 000 × g for 5 minutes at 4 °C. The resulting chromatin-containing supernatant was diluted 1:10 with ChIP dilution buffer (1% Triton X-100, 1.2 mM EDTA, 16.7 mM Tris-HCl pH 8, 167 mM NaCl, and 1× EDTA-free protease inhibitor cocktail (Roche #CO-RO)). Diluted chromatin was distributed into DNA LoBind tubes (Eppendorf) and incubated overnight at 4 °C with a 1:500(v/v) antibody on a rotator set at 10 rpm. Antibodies were sourced from Cell Signaling Technology (anti-H3: #4620, H3K4me3: #9751S, H3K9ac: #9649S, H3K79me2: D15E8,

H4K20me3: #5737S) and Abcam (H3K36me3: ab9050). Immunoprecipitation was carried out using a 1:1 mixture of protein A and G Dynabeads (Thermo Fisher Scientific #10004D and #10002D). Following binding and sequential wash steps, immune complexes were eluted in 100 μl of Direct Elution Buffer (0.5% SDS, 5 mM EDTA, 10 mM Tris-HCl pH 8, 300 mM NaCl). Cross-link reversal was achieved by incubating samples at 65 °C overnight with intermittent shaking. DNA was purified following digestion with Proteinase K (Fisher Scientific #11826724) and RNase A (Roche #10109142001). DNA extraction was performed using phenol/chloroform/isoamyl alcohol (25:24:1), followed by centrifugation at 13 800 × g for 15 minutes at 4 °C. The aqueous phase was transferred to fresh DNA low binding tubes, mixed with 1.25 ml of 100% ethanol, 50 µl of 3 M sodium acetate (pH 5.2), and 4 µl of glycogen (20 mg/ml), and incubated at –80 °C for at least 1 hour (or overnight) for DNA precipitation. DNA was pelleted by centrifugation at 13 800 × g for 15 minutes at 4 °C, washed with 70% ethanol, and centrifuged again under the same conditions. Pellets were air-dried and resuspended in 0.1× TE buffer. Library preparation was conducted using the NEBNext® Ultra™ II DNA Library Prep Kit (New England Biolabs #E7645S), and sequencing was carried out on the Illumina HiSeq 3000 platform, targeting 20 millions of 150-bp paired-end reads per sample.

To process the ChIP-seq data, we used nf-core/chipseq v2.0.0 with default options ^50^. Publicly available datasets from wild-type male and female gametophytes of *Ectocarpus*^8^ were retrieved and processed using the same workflow for consistency. Biological replicates for each species were aligned to their corresponding reference genome (see **Table S1**). Replicates showing high correlation, as determined by Spearman’s coefficient using `multiBamSummary` and `plotCorrelation` from deepTools v3.5.1^51^ were merged using samtools merge for downstream analyses. Peaks were called with `macs2 (defaul parameters)`.

Normalised log₂ coverage tracks, relative to total H3, were generated using `deepTools bamCompare` with a bin size of 10 bp, `--scaleFactorsMethod readCount`, and `- -operation log2`. Using the deepTools, we computed the correlation matrices via `multiBigwigSummary` (in bins mode) followed by `plotCorrelation`. Genome-wide signal profiles were visualized in IGV v2.18.4.

### RNA-seq

RNA-seq data was generated from culture with same conditions to match the histone PTM data with the gene expression data. Each RNA-seq were carried out in triplicates. For each replicate, approximately 10 mg of algal tissue was gently blotted dry and immediately flash-frozen in liquid nitrogen. Total RNA was extracted following the method described by^10^. Briefly, frozen tissue was ground in liquid nitrogen and incubated at 65 °C in 700 μL of preheated CTAB3 extraction buffer (100 mM Tris-HCl pH 8, 1.4 M NaCl, 20 mM EDTA pH 8, 2% CTAB, 2% PVP, and 1% β-mercapntoethanol). The lysate was vortexed and maintained at 65 °C for 5–20 minutes. Phase separation was achieved by extraction with chloroform:isoamyl alcohol (24:1), followed by two rounds of centrifugation at 10 000 × g for 15 minutes at 4 °C. RNA was precipitated overnight at –20 °C using 3 M LiCl and 1% β-mercaptoethanol, pelleted by centrifugation at 10 000 × g for 1 hour at 4 °C, washed with cold 70% ethanol, and resuspended in RNase-free water. Genomic DNA contamination was removed using the TURBO DNase Kit (Thermo Fisher, AM1907). RNA-seq libraries were prepared using the NEBNext® Ultra™ II Directional RNA Library Prep Kit (New England Biolabs, E7760S) and sequenced on the Illumina Next Seq 2000 platform, generating 25–30 million 150-bp paired-end reads per sample.

Reads were processed using the nf-core/rnaseq pipeline v3.12.0 ^50^. Genome assembly versions are described in **Table S1**. Publicly available RNA-seq datasets from *Ectocarpus sp.7* male and female gametophytes^8^ were reprocessed using the same workflow for consistency. Reproducibility across biological replicates was confirmed via Spearman correlation. Differential expression analysis between male and female was performed with DESeq2 (v1.42.1)^52^, identifying differentially expressed genes (DEGs) based on a |log₂ fold change| ≥ 1 and an adjusted p-value < 0.05.

### Generation of histone mark profiles

To investigate the distribution of histone modifications across genes grouped by expression level, we used a custom bash script to automated signal processing and plotting using with deepTools v3.5.1 within a conda environment. Genes were divided into five groups: one group comprising genes with zero expression (0 TPM), and the remaining genes divided into four groups according to expression level quartiles one where there were no expression and the rest split by quartiles (calculated for each sample). Gene coordinates grouped by expression quantiles were provided as BED files, and ChIP-seq signal was taken from precomputed bigWig tracks. Signal matrices were generated using `computeMatrix scale-regions -a 1000 -b 1000 -bs 100 —skipZeros`, which summarized ChIP-seq signal over gene bodies (represented as 5 kb) and 1 kb flanking regions, with a bin size of 100 bp. Regions with zero signal were excluded. The resulting matrices were visualized using `plotProfile--plotType se`, producing average signal profiles per quantile group.

### Chromatin emission state and signature inference

#### Detection of emission states with hiHMM

The five chromatin marks were analysed using hiHMM ^21^ to annotate each genome with emission states. Input files for hiHMM were generated in several steps following the recommendations from ^21^. First, BedGraph files were produced using `bamCompare -bs 200 --scaleFactorsMethod readCount - -pseudocount 0.5 --operation log2 -of “bedgraph”`. Then BedGraph files were normalized prior to modeling using a short Python script that applied `sklearn.preprocessing.StandardScaler.fit_transform()`to the signal column (the 4th). Quality check involving visual inspection of quantile-quantile plots confirmed standardization. Normalized BedGraphs were reformatted into chromosome-wise matrices with genomic bins as rows and samples as columns and file and chromosome names were standardized as required.

The hiHMM model was trained on a reduced subset of chromosomes (one autosome per sex, plus one male and one female sex chromosome or sex homolog when available), which reduces computational complexity without affecting model quality according to the authors^21^. For each species, we selected the two longest scaffolds together with the available sex chromosome(s). The list of training chromosomes and run parameters are provided in **Table S11**. The optimised model initiates with K_0_ = 7 and ends up with K = 27 states plus one “E0” state containing regions where no reads were assigned. The decoding part was carried on all scaffolds. The decoding part was carried on all scaffolds. The emission matrix can be found in **Fig. 2B** and transition matrix can be found in **Fig. S13** as well as the input files used in **Supplemental Dataset 1**. Model 2 was favored, as Model 1 did not provide an integrated or streamlined representation of the data. To examine the hiHMM model, we adapted the advanced functions of ChromHMM^53^. The optimization of hiHMM model was obtained using `ChromHMM CompareModels` function where hiHMM model was tested for different value of K_0_. From these comparisons, we decided to use `K0 = 7` (as the default). The overlap enrichment of genomic features was produced with `ChromHMM OverlapEnrichment`.

#### Definition of chromatin signatures

To streamline the analysis, emission states assigned to each gene were consolidated into five broader categories based on their predominant emission enrichment with genomic features as done in^54^. States showing strong enrichment at transcription start sites (TSS) were classified under the “TSS” category (E5, E10, E12). Emission states predominantly associated with gene bodies were grouped into the “Genic” category (E8, E2, E7, E9, E25). States displaying mixed enrichment patterns—with or without H3K36me3—were collectively grouped under “mixed,” due to the absence of clear feature-specific enrichment (E14, E17, E19, E4, E24, E18, E15, E11, E6, E20, E26, E22, E23). A distinct “Silenced” category was defined for states marked exclusively by H3K79me2 and H4K20me3 (specifically E27 and E1). Finally, states characterized by minimal or no detectable histone modification signals were assigned to the “Null states” group (E0, E3, E13, E16, E21). Each gene was then annotated based on its combination of these five chromatin categories, resulting in a defined set of unique chromatin profiles referred to as ‘chromatin signatures’.

Genes were assigned to the null signature group (S1) if they were associated solely with low-signal states (E0, E3, E13, E16, E21), and no other chromatin category. In total, this classification yielded 16 distinct chromatin signatures (S1–S16) across the six analysed genomes. Signatures were then labeled following the increasing gene expression median across all species.

Chromatin signatures of genes were extracted from **Table S5-10** and compared between samples. Class transitions were summarized with sankey plots using `ggalluvial::geom_alluvium()` to visualise flux as well as stacked bar plot using `ggplot2::geom_col()` in R. Enrichment of chromatin changes was examined by different category type such as gene location (autosomes, PAR, SDR, sex-homolog chromosome) and expression bias (female-biased, male-biased, unbiased). Statistical relevance was assessed with chi-square test using `stats::chisq.test()` in R, comparing the count of one category against the total genes. P-values were Bonferroni adjusted using `p.adjust(method = “bonferroni”)`.

### Evolutionary analysis of chromatin signatures

We analysed the overall conservation of chromatin signatures by counting the number of one-to-one orthologs with the same chromatin signature label across all six species. To examine the significance of the observed value, we performed a permutation test by reshuffling the chromatin signature labels across orthologs for each species and recomputing the number of overlaps. The permuted values represent the null assumption of no conservation of chromatin signatures. The same procedure was followed for pairwise comparisons (against *Ectocarpus*). These analyses were enabled by rsample v1.2.1 (https://rsample.tidymodels.org). Female chromatin signature data was used for dioicous species in this and following analyses. *P*-values were computed from the 10,000 permutation results by comparing the observed statistic to the empirical null distribution.

To examine the influence of sex chromosomes on the signature conservation, we calculated the percentage of observed cross-species pairwise overlaps separately for autosomes and sex chromosomes for each focal species. We only considered the chromosome type in the focal species and not the target species of the comparison due to the high evolutionary turnover rate of one-to-one orthologues in the sex chromosome. This analysis included the former sex chromosome of *D. dudresnayi* but not *S. ischiensis* since the sexual system is unknown. Statistical comparisons between autosomes and sex chromosomes were conducted using paired, one-tailed Wilcoxon rank sum tests.

To profile the gene-wise evolutionary signal associated with each chromatin signature, we performed genomic phylostratigraphy using GenEra^23^ to infer the evolutionary age of each gene, with the resulting phylostratigraphic rank (phylorank) ranging from 1 (cellular organisms) to 15 (species-level) in *Ectocarpus*. It should be noted that the phylorank of the youngest genes differ between species due to differences in the presence of genomic data at each taxonomic node. For the orphan gene analysis, gene models with the highest phylorank for each species were considered as orphan genes, which we then obtained their prevalence (in %) in each chromatin signature.

Alongside the evolutionary signals, we profiled the expression variability associated with each chromatin signature. Using a recently published developmental RNA-seq dataset of *Ectocarpus*^55^ we calculated *tau* scores^56^ to estimate the expression specificity of genes, which ranges from 0 (broadly-expressed) to 1 (developmental stage-specific expression), using the median expression values for each gene across replicates. For other species, which lack comparably comprehensive RNA-seq datasets, we calculated expression variability using the RNA-seq data produced in this study as input. Specifically, we computed the relative entropy (using `KL.empirical()` from entropy v1.3.2) for each gene between the observed log₂(TPM+1) expression values across replicates and a null assumption of uniform expression for each gene. To reduce the effect of noise in lowly expressed genes, genes with mean TPM below 3 were discarded for this analysis.

### Data analytics and graphics

Statistical analysis, data processing and visualisation were done using R v4.3.3 with the following packages: tidyverse v2.0.0^57^ (ggplot2 v3.5.2, dplyr v1.1.4, tidyr v1.3.1, tibble v3.3.0, stringr v1.5.1, readxl v1.4.3), rtracklayer v1.64.0^58^, duckplyr v0.4.1 (https://duckplyr.tidyverse.org/), GenomicRanges v1.56.2, ggalluvial v0.12.5 (http://corybrunson.github.io/ggalluvial/), ggrastr v1.0.2 (https://github.com/VPetukhov/ggrastr), and rsample v1.2.1 (https://rsample.tidymodels.org). Statistical tests and significance levels are indicated in the text, figure legends and **Table S12**. Scripts are provided at https://github.com/jerovign/PhaeoChromo.

### Data availability

Mass spectrometry data are available via ProteomeXchange under identifier PXD065559. ChIP-seq and RNA-seq short-read data has been uploaded to NCBI under BioProject PRJNA1328953. Supplemental Dataset 1 can be accessed at https://doi.org/10.17617/3.TDGYHS.

## Contributions

SMC, MB conceived and designed the experiments. JV performed experiments and the algae culture was supported by RL. JV, JSL, MB, FH analyzed the data. PL provided unpublished data and performed experiments. Project Administration: SMC and MB; Resources: SMC; BL carried out the MS experimental work. DL supervised MS data analysis. SMC and MB wrote the manuscript with input from JV and JSL. All authors read and approved the manuscript.

## Acknowledgements

This study was funded by the Max Planck Society, the European Research Council grant 864038 (SMC), the Bettencourt Foundation (SMC) and the Moore Foundation (SMC). The MS platform at the Curie Institut is supported by Région Ile-de-France (N°EX061034) and ITMO Cancer of Aviesan and INCa supported by funds administered by INSERM (no. 21CQ016-00). The LSMP thanks Patrick Poullet from the bioinformatics Core facility (CUBIC) of the Institut Curie U1331 for the continuous development of myProMS. JV and JSL are thankful to the IMPRS ‘From molecules to organisms’.

## Supplemental Figures

**Fig. S1:**
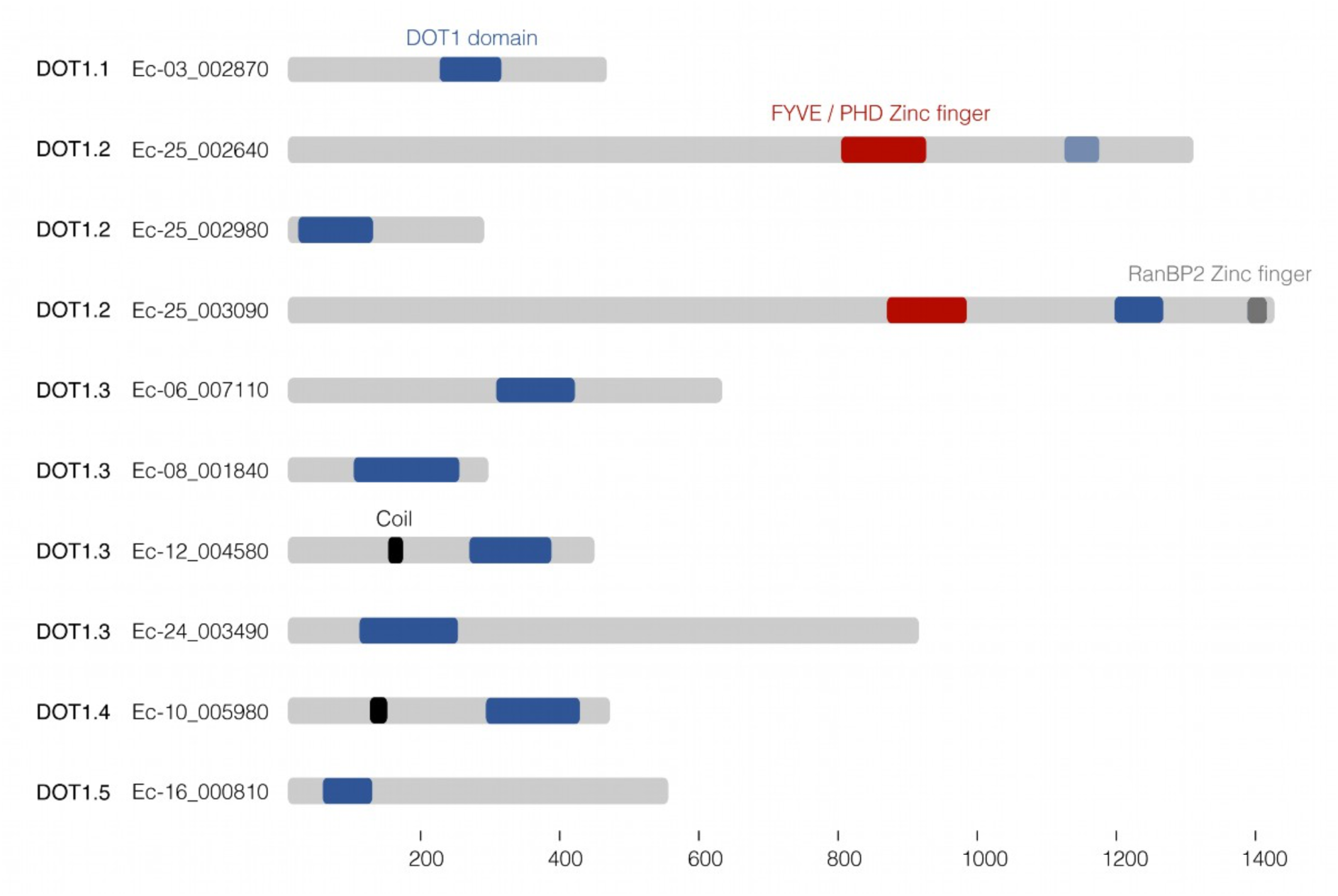
*Ectocarpus* encodes ten DOT1-domain containing proteins that form part of five distinct gene families.

**Fig. S2:**
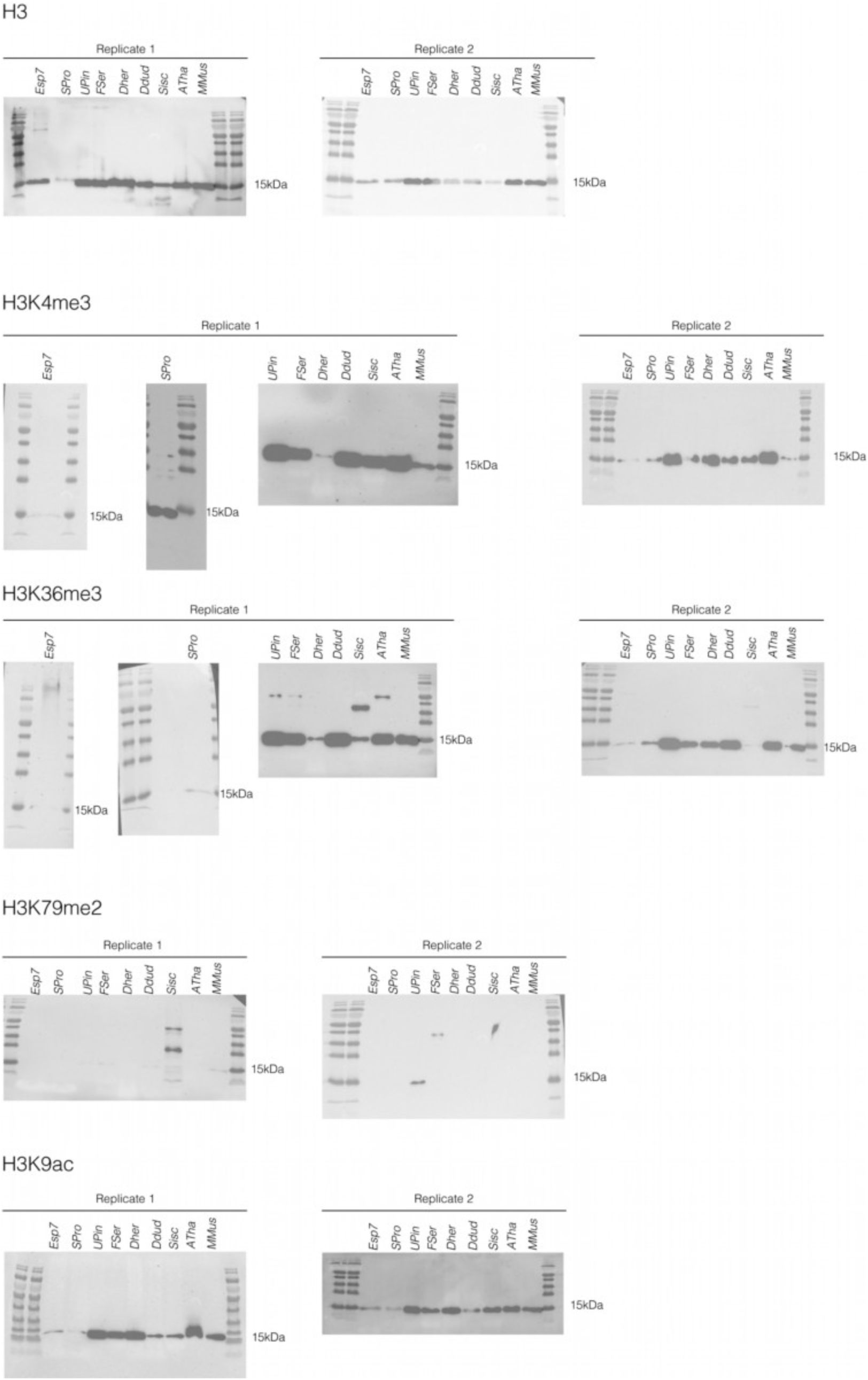
Uncropped Western blots of selected hPTMs across species and model organisms such as *Arabidopsis thaliana* (*ATha*) and *Mus musculus* (*MMus*).

**Fig. S3:**
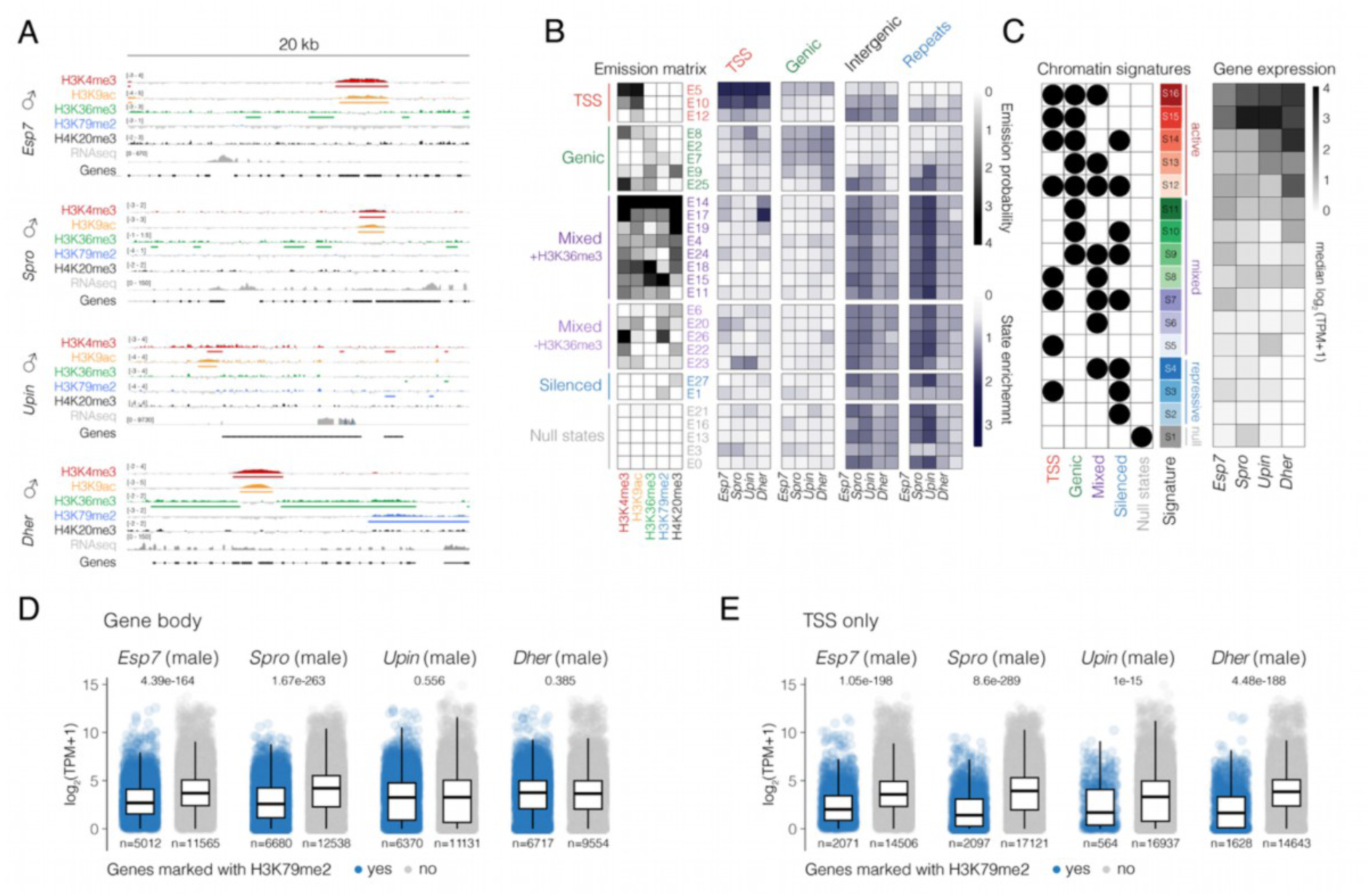
(**A**) Representative genome browser tracks of ChIP-seq and RNA-seq datasets in male datasets of the dioicous species. ChIP-seq coverage is represented as the log_2_ ratio of IP DNA relative to histone H3, with the range indicated on each track. ChIP-seq peaks for each hPTM are indicated under their respective track. (**B**) Enrichment of each chromatin state in genomic features in the male samples of the four dioicous species. (**C**) Median RNA-seq expression level (log_2_ TPM+1) of genes associated with each signature in the male samples of the four dioicous species. (**D-E**) Gene expression in males for genes with (blue) or without (grey) an H3K79me2 peak overlapping their gene body (**D**) or TSS (**E**).

**Fig. S4:**
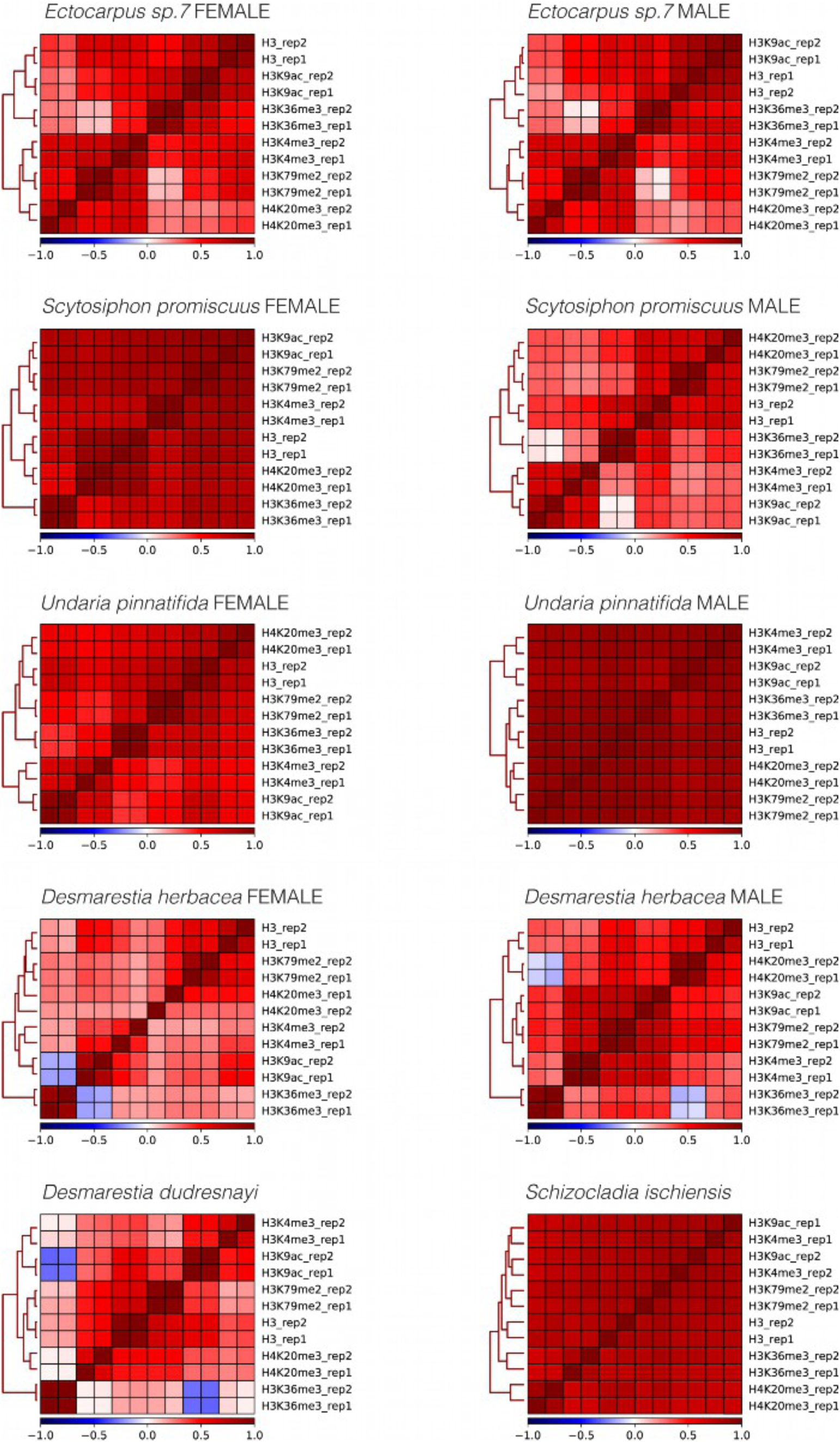
Spearman correlation matrix of the ChIP-seq datasets for each species.

**Fig. S5:**
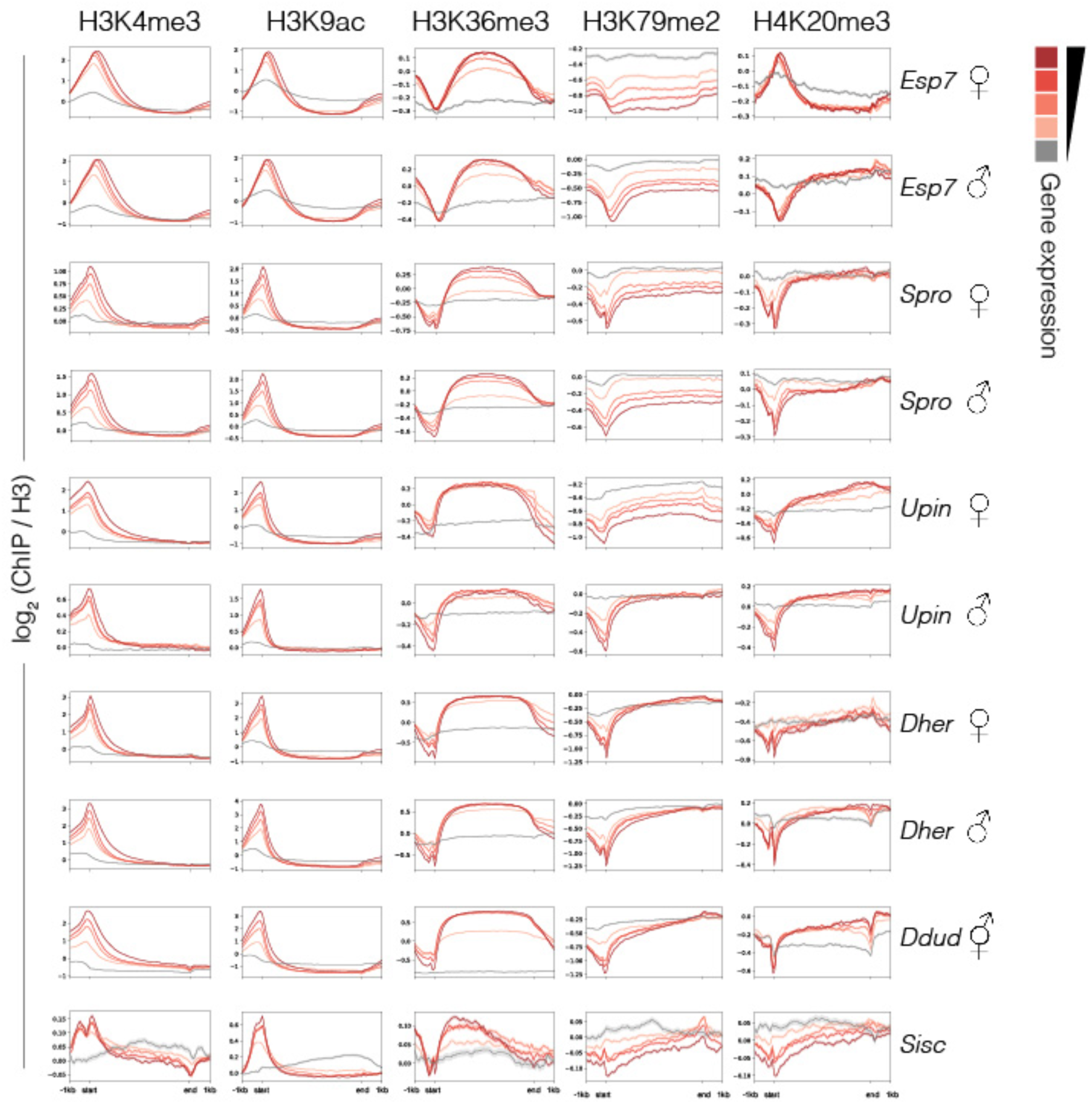
Metaplots of H3K4me3, H3K9ac, H3K36me3, H3K79me2 and H4K20me3 coverage over genes grouped by gene expression levels in each sample. Gene bodies are plotted as proportional lengths, upstream and downstream intergenic regions in kilobases.

**Fig. S6:**
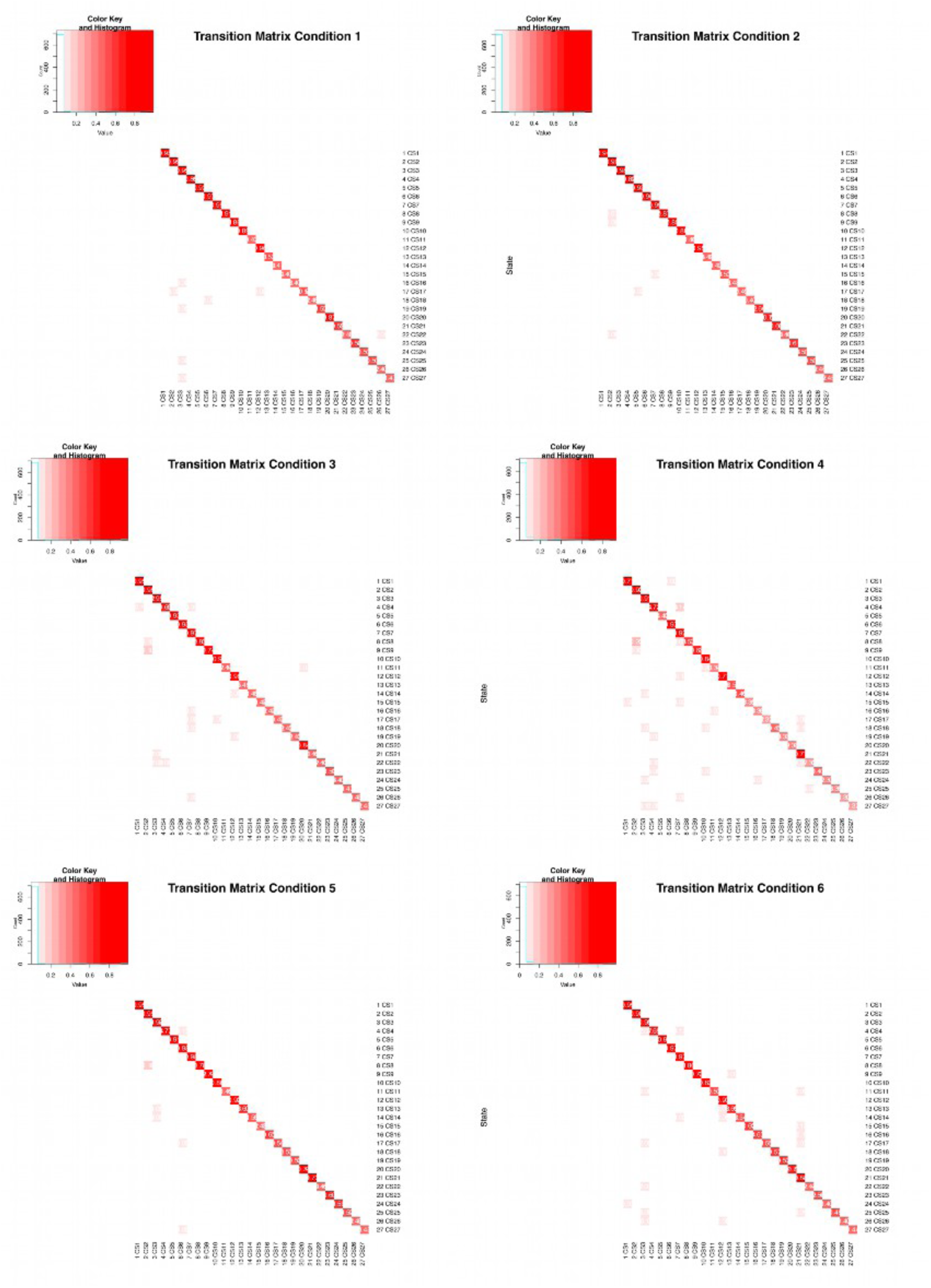
Transition matrices of hiHMM model.

**Fig. S7:**
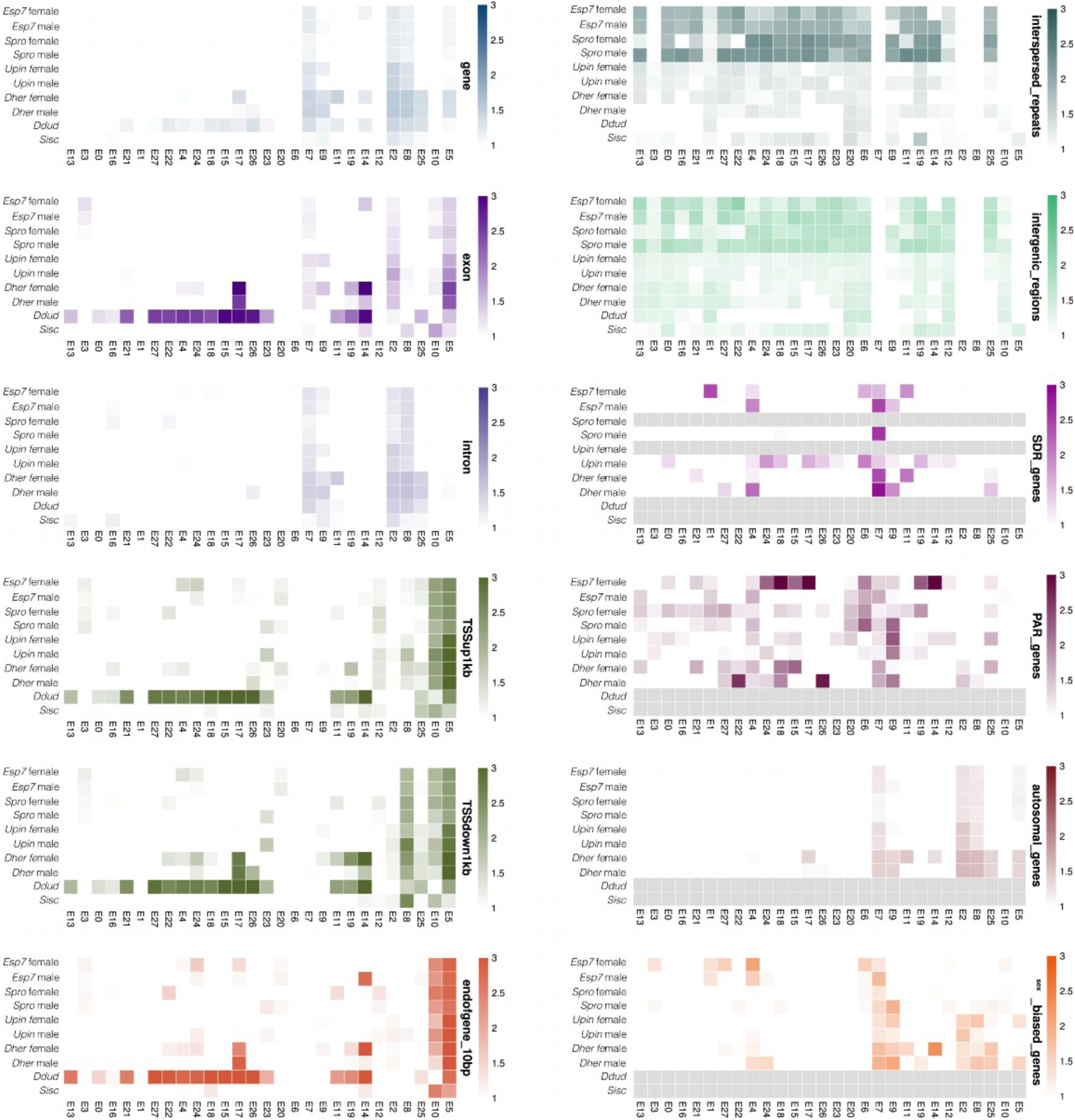
Enrichment of hiHMM emission states over genomic features.

**Fig. S8:**
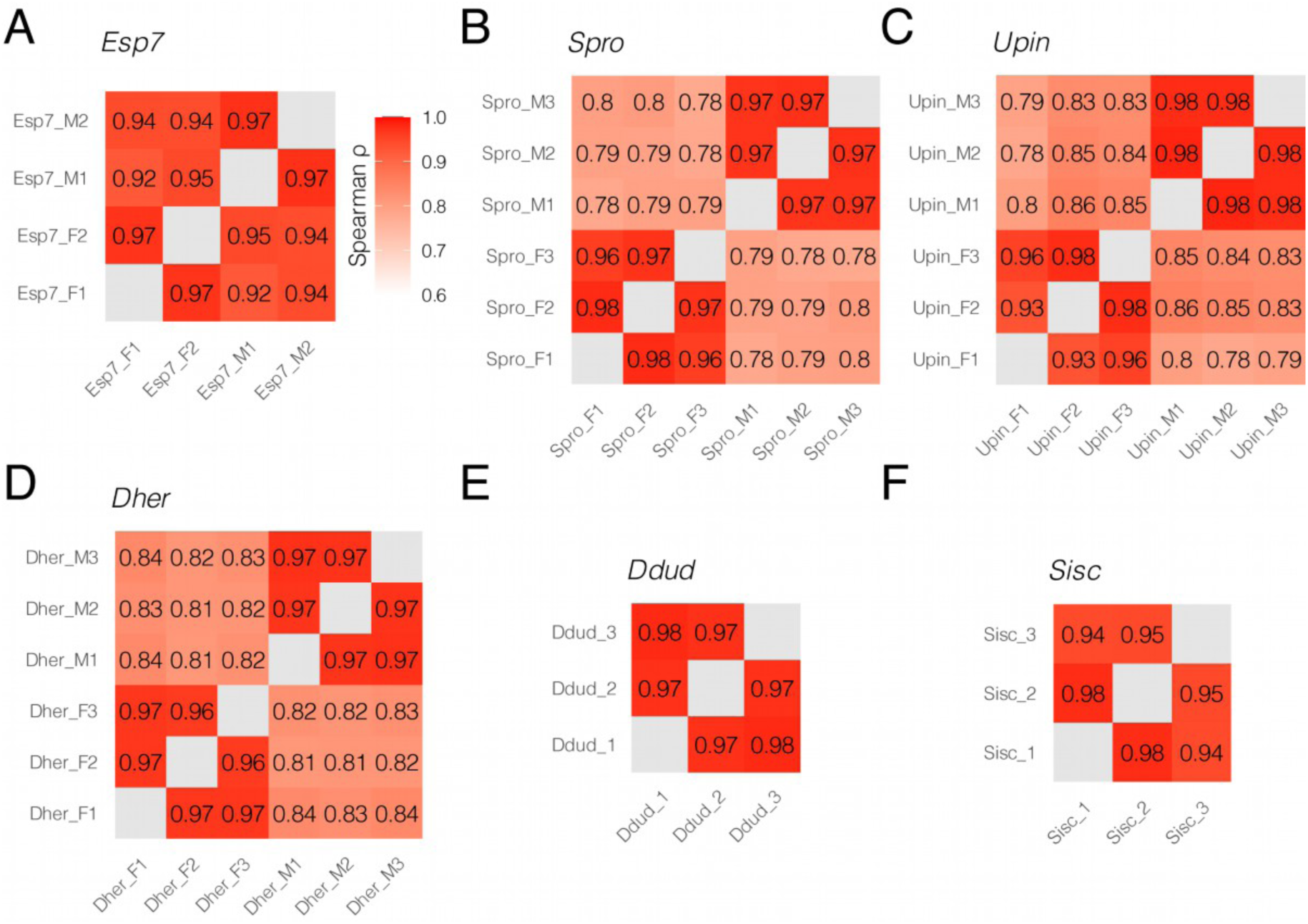
Cross-correlation matrices for the RNAseq datasets in this study. (**A**) *Ectocarpus* (from^8^ PRJNA671807), (**B**) *Scytosiphon promiscuus*, (**C**) *Undaria pinnatifida*, (**D**) *Desmarestia herbacea*, (**E**) *Desmarestia dudresnayi*, and (**F**) *Schizocladia ischiensis*. Male and female samples are indicated by suffix “_M” or “_F”, respectively. Correlation is measured using Spearman’s ρ.

**Fig. S9:**
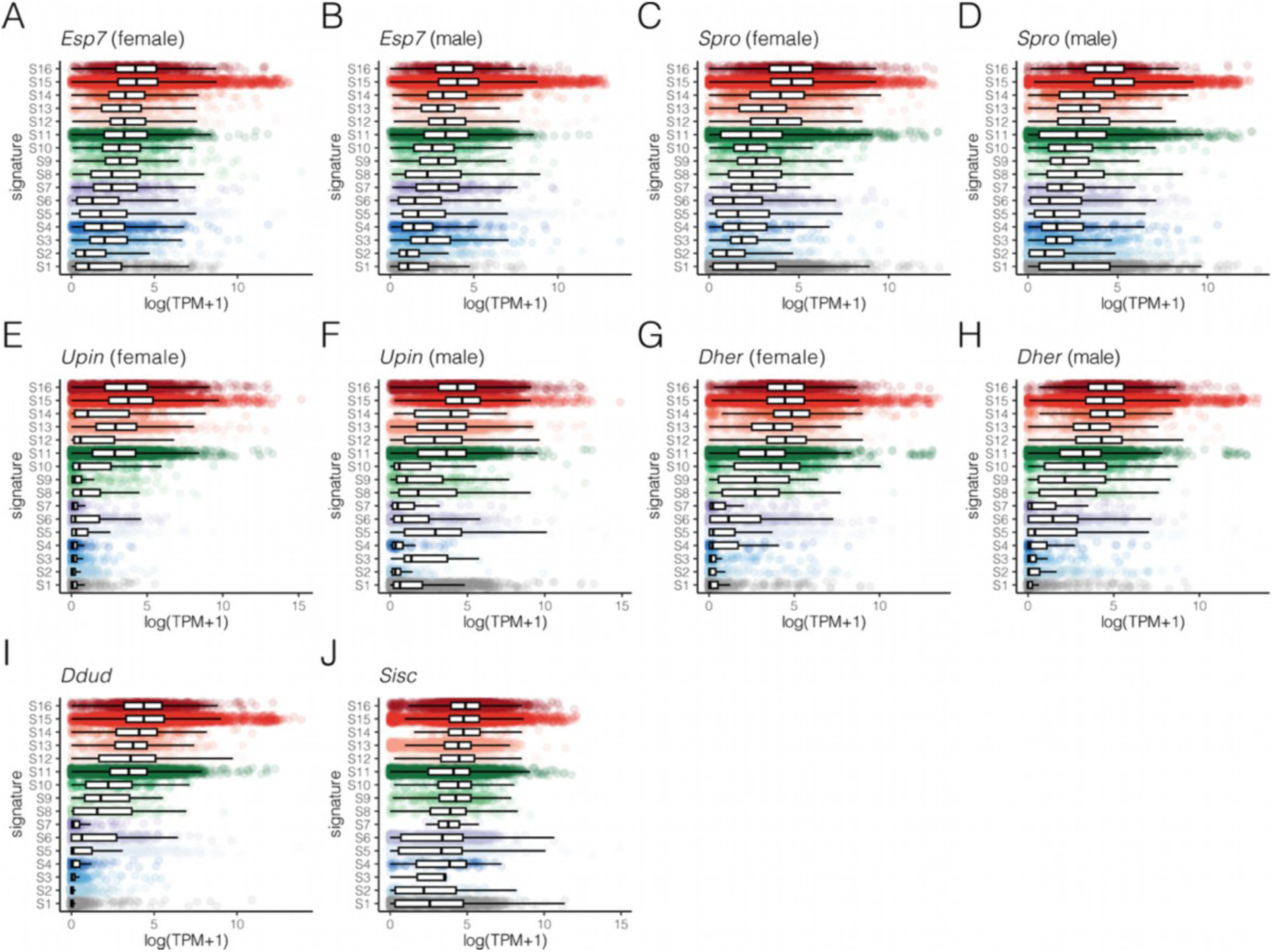
Gene expression levels of genes group by their assigned chromatin signature across all ChIP-seq datasets generated in this study. (**A**) *Ectocarpus* sp 7 female, (**B**) *Ectocarpus* sp 7 male, (**C**) *Scytosiphon promiscuus* female, (**D**) *S. promiscuus* male, (**E**) *Undaria pinnatifida* female, (**F**) *U. pinnatifida* male, (**G**) *Desmarestia herbacea* female, (**H**) *D. herbacea* (male), (**I**) *Desmarestia dudresnayi*, and (**J**) *Schizocladia ischiensis*.

**Fig. S10:**
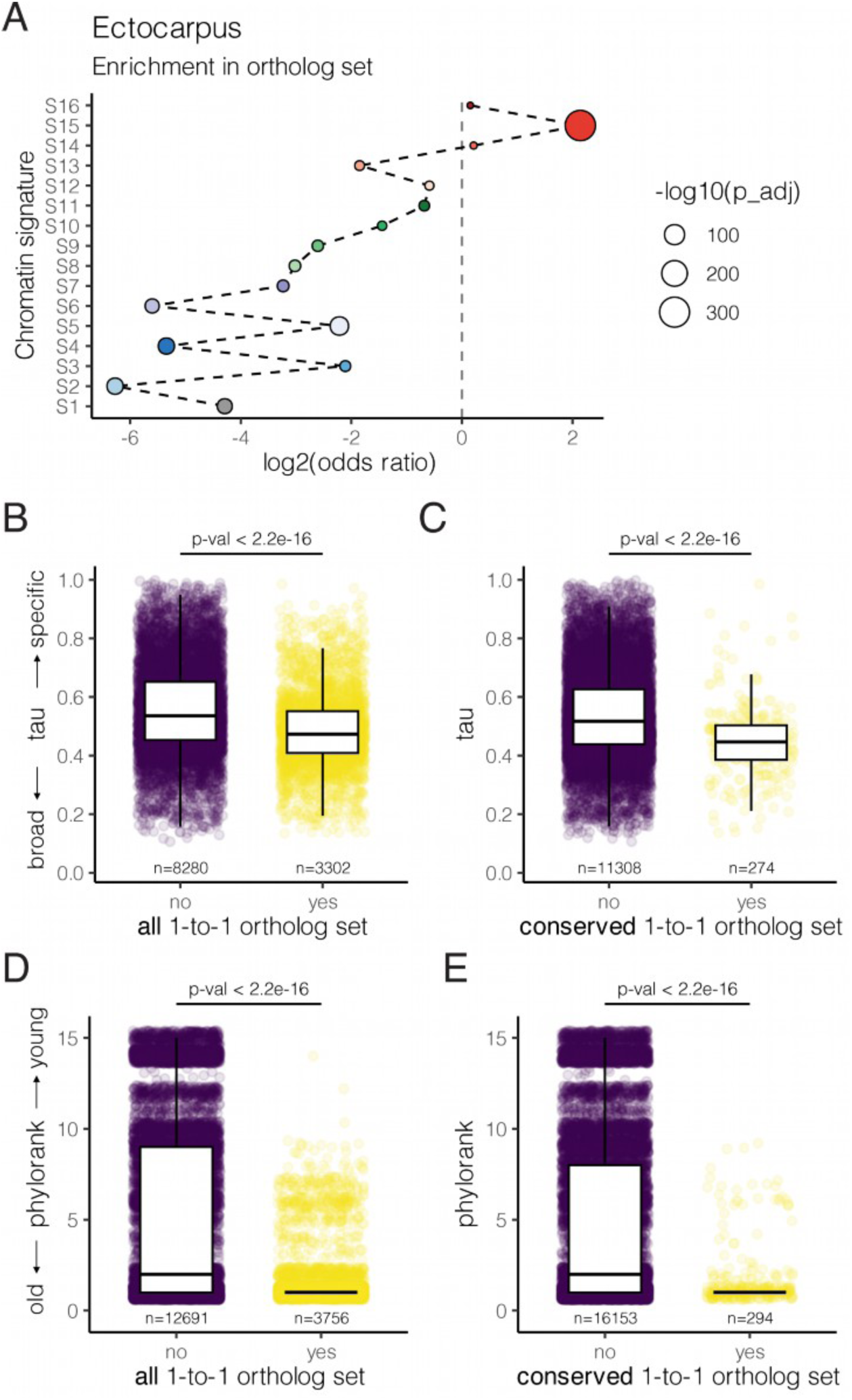
Expression and evolutionary profile of genes in the one-to-one ortholog set compared to all other genes in *Ectocarpus*. (**A**) Chromatin signatures enriched in the one-to-one ortholog set. The Benjamini-Hochberg Procedure was used to adjust the p-values computed from Fisher’s exact test. (**B**) Distribution of expression specificity score in all genes on the one-to-one ortholog set and (**C**) in a subset of one-to-one orthologs with conserved chromatin signature across all species. (**D**) Distribution of gene age (phylorank) in all genes on the one-to-one ortholog set and (**E**) in a subset of one-to-one orthologs with conserved chromatin signature across all species. Related to Fig. 3A. The Kruskal–Wallis test is used to compute the p-values in (**B**-**E**).

**Fig. S11:**
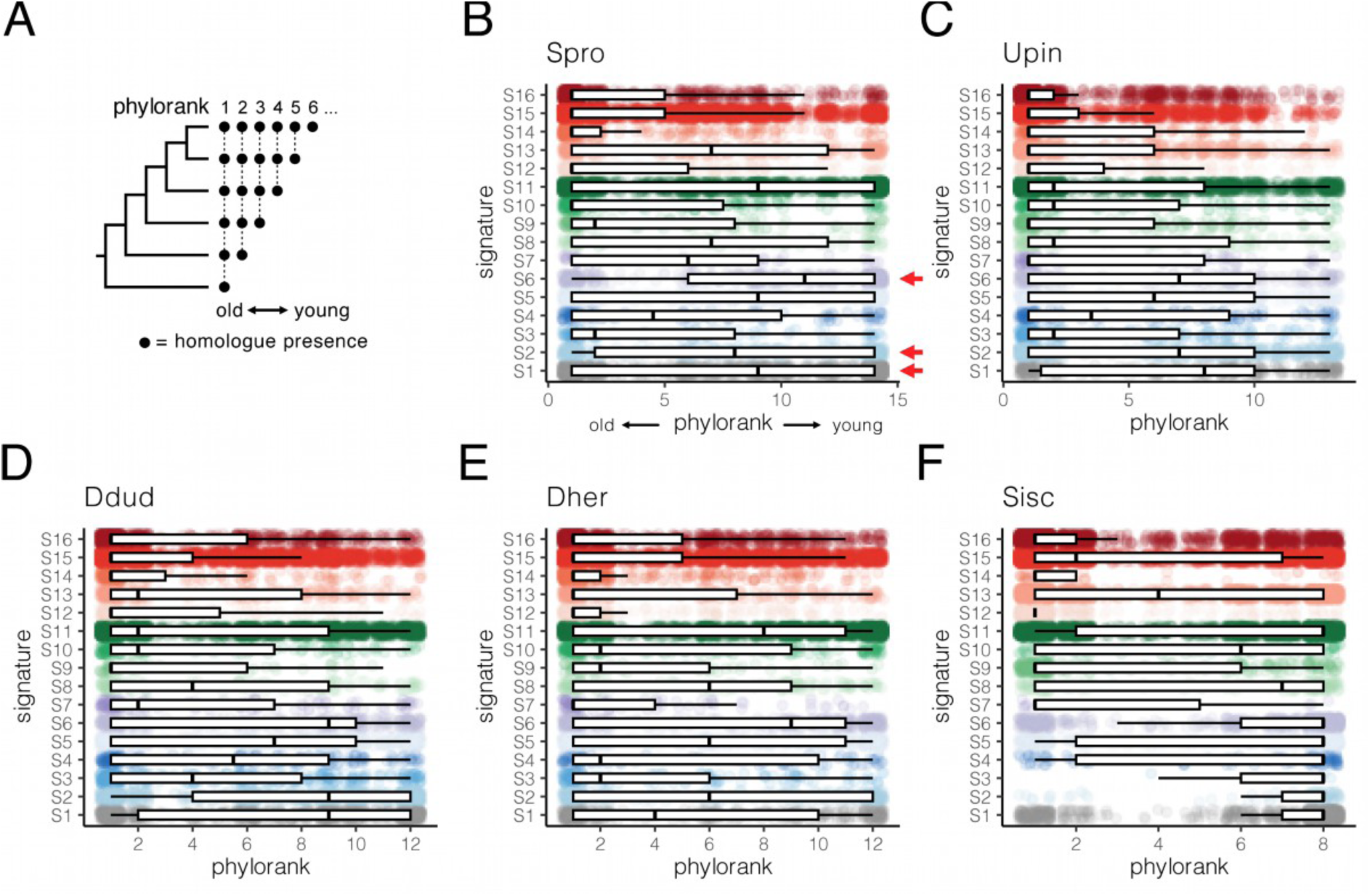
Distribution of gene age across each chromatin signature. (**A**) Summary of the approach to infer gene age (phylorank) via pairwise sequence alignment. The resulting gene age distribution in each chromatin signature for (**B**) *Scytosiphon promiscuus*, (**C**) *Undaria pinnatifida*, (**D**) *Desmarestia dudresnayi*, (**E**) *Desmarestia herbacea*, and (**F**) *Schizocladia ischiensis*. The red arrows mark the chromatin signatures with consistently high distribution of young genes across species. Related to Fig. 3C. Female chromatin signatures were used for dioicous species.

**Fig. S12:**
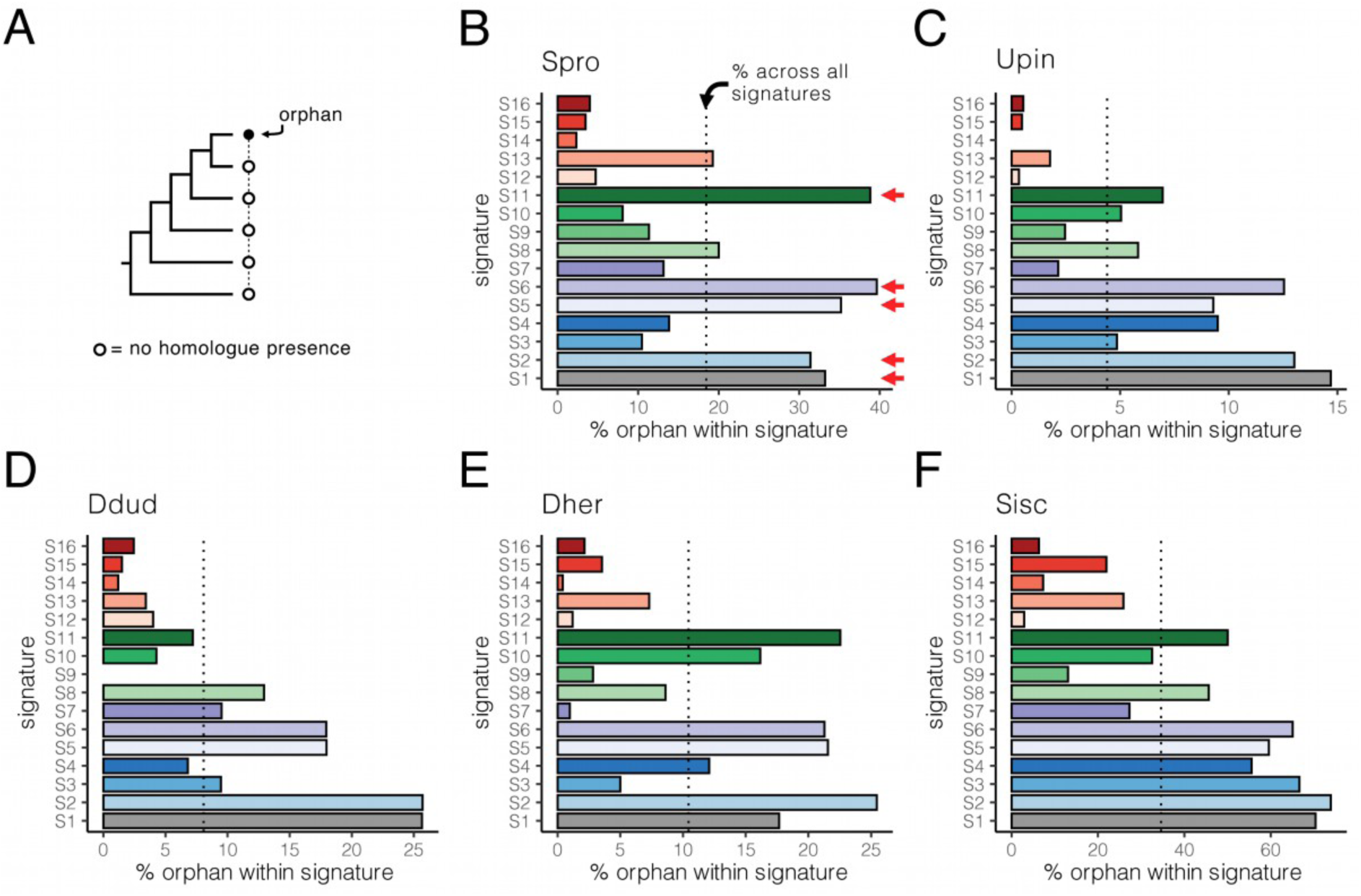
Percentage of orphan genes assigned to each chromatin signature. (**A**) Summary of the approach to infer orphan genes via pairwise sequence alignment. The resulting percentage of orphan genes detected in each chromatin signature for (**B**) *Scytosiphon promiscuus*, (**C**) *Undaria pinnatifida*, (**D**) *Desmarestia dudresnayi*, (**E**) *Desmarestia herbacea*, and (**F**) *Schizocladia ischiensis*. The red arrows mark the chromatin signatures with consistently high orphan gene presence across species. Related to Fig. 3D. Female chromatin signatures were used for dioicous species.

**Fig. S13:**
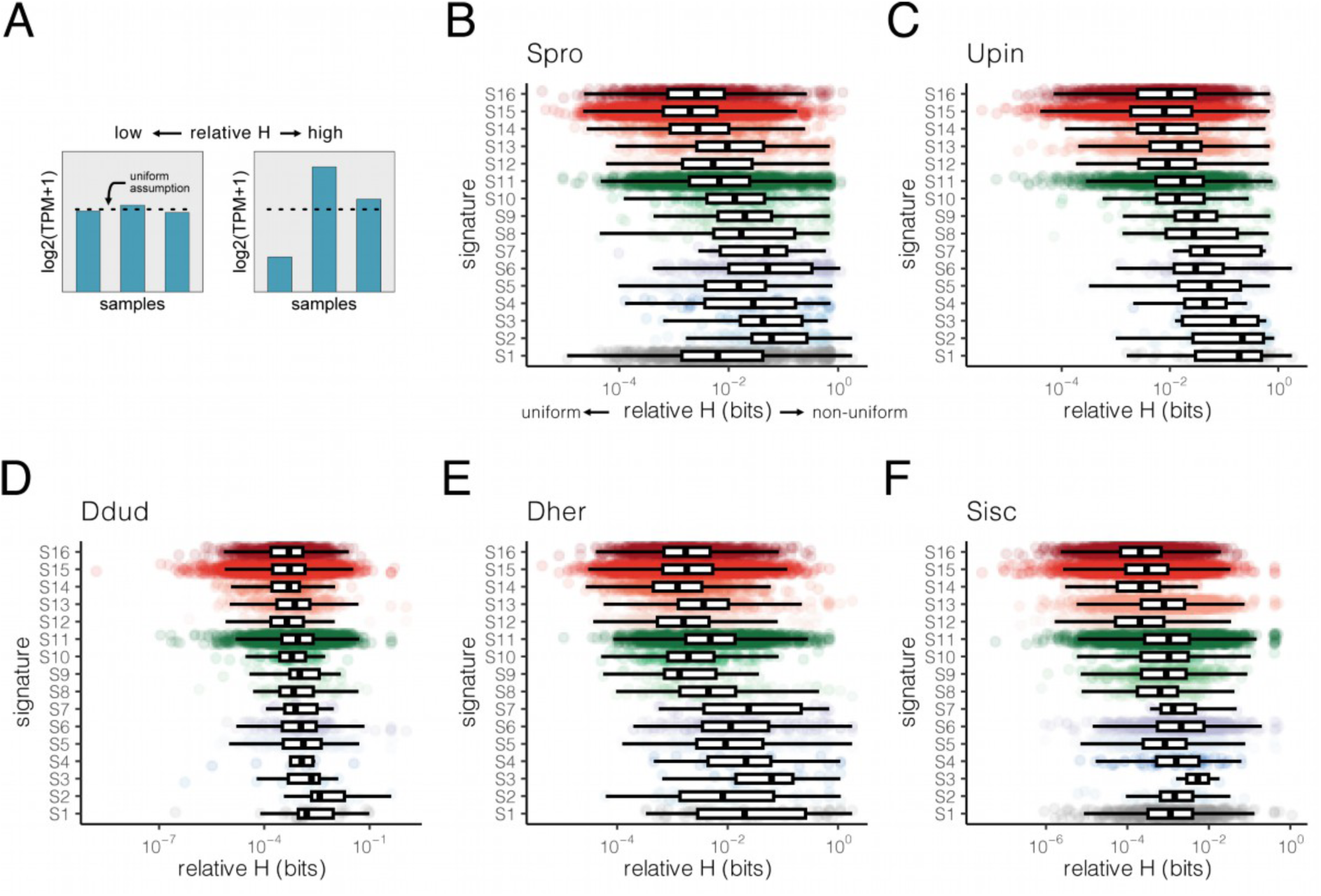
Distribution of expression uniformity across RNA-seq libraries for genes assigned to each chromatin signature. (**A**) Summary of the relative entropy (relative H) approach to compare the observed log₂(TPM+1) expression values across replicates and a gene-specific null assumption of uniform expression values. The resulting relative H scores across chromatin signature for (**B**) *Scytosiphon promiscuus*, (**C**) *Undaria pinnatifida*, (**D**) *Desmarestia dudresnayi*, (**E**) *Desmarestia herbacea*, and (**F**) *Schizocladia ischiensis*. Related to Fig. 3E.

**Fig. S14:**
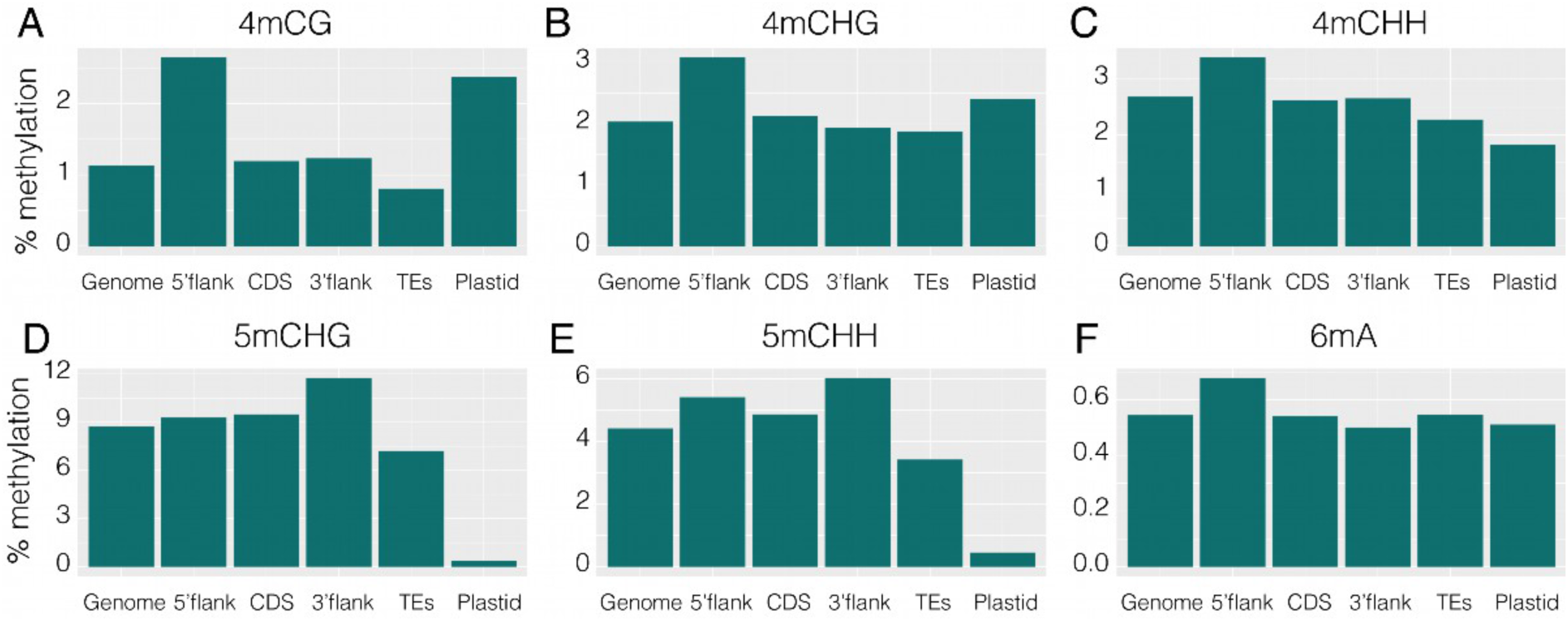
Average genome-wide levels of (**A-**C) 4mC, (**D-**E) 5mC and (**F**) 6mA in all relevant contexts at different genomic features of the nuclear genome and plastid genome in *S. ischiensis*.

## Supplemental Table Legends

Table S1 – Species used, strain and genome assembly reference.

Table S2 – Summary of mass spectrometry results.0

Table S3 - Quality of the protein samples for mass spectrometry analysis.

Table S4 - ChIP-seq quality metrics.

Table S5 - Data table for *Ectocarpus* genes.

Table S6 - Data table for *Scytosiphon promiscuus* genes.

Table S7 - Data table for *Undaria pinnatifida* genes.

Table S8 - Data table for *Desmarestia herbacea* genes.

Table S9 - Data table for *Desmarestia dudresnayi* genes.

Table S10 - Data table for *Schizocladia ischiensis* genes.

Table S11 - Orthogroups across species.

Table S12 - Enrichment and statistics of signature switches between sexes in brown algae.

Table S13 - List of chromosomes used to train hiHMM model and specific command lines.

